# Improved analysis of (e)CLIP data with RCRUNCH yields a compendium of RNA-binding protein binding sites and motifs

**DOI:** 10.1101/2022.07.06.498949

**Authors:** Maria Katsantoni, Erik van Nimwegen, Mihaela Zavolan

## Abstract

We present RCRUNCH, an end-to-end solution to CLIP data analysis for identification of binding sites and sequence specificity of RNA-binding proteins. RCRUNCH can analyze not only reads that map uniquely to the genome, but also those that map to multiple genome locations or across splice boundaries, and can consider various types of background in the estimation of read enrichment. By applying RCRUNCH to the eCLIP data from the ENCODE project, we have constructed a comprehensive and homogeneous resource of *in vivo*-bound RBP sequence motifs. RCRUNCH automates the reproducible analysis of CLIP data, enabling studies of post-transcriptional control of gene expression.

## Background

Throughout their life cycle, from transcription to maturation, function and decay, RNAs associate with RNA-binding proteins (RBPs) to form ribonucleoprotein complexes (RNPs) or higher-order RNA granules (e.g paraspeckles, Cajal bodies) [1]. RBPs are abundant in prokaryotes as well as eukaryotes, and methods such as RNA interactome capture (RIC) [2,3] revealed that over a thousand human and mouse proteins have RNA-binding activity. An RBP is typically composed of multiple RNA-binding domains (RBDs) coming from a limited repertoire [4], and binds to a specific sequence motif and/or secondary structure element. The functional diversity of RBPs rests on the number and arrangement of RBDs that they contain [5], though methods like RIC have uncovered many proteins that have RNA-binding activity, despite lacking a known RBD [2].

As RBPs participate in all steps of RNA metabolism, it is not surprising that they have been implicated in many diseases [6,7]. However, the critical targets in a particular context are often unknown. The method of choice for mapping the binding sites of an RBP *in vivo* and transcriptome-wide is crosslinking and immunoprecipitation (CLIP). Introduced in the early 2000s [8], CLIP has a number of variants, all exploiting the photoreactivity of nucleic acids and proteins. Briefly, ultraviolet light is used to crosslink RBPs to RNAs, the regions of the RNAs that are not protected by RBPs are enzymatically digested, the RBP of interest is purified along with the crosslinked RNAs, and finally the purified RNA fragments are reverse-transcribed and sequenced. One of the main differences between CLIP variants is in the nature of the cDNAs that end up being sequenced. These can either be the result of aborted reverse transcription at the crosslinked site [9], where a bulky adduct remains after protein digestion, or the result of reverse transcription through the site of crosslink, which often results in characteristic mutations in the cDNAs [10,11]. Although the general expectation is that extensive purification leads to a relatively pure population of target sites for a given protein, inspection of the genome coverage by sequenced reads indicates substantial non-specific background. Various approaches have been proposed for background correction, but a systematic benchmarking of these approaches is still lacking (discussed in [12]).

CLIP is analogous to chromatin immunoprecipitation (ChIP), a technique that has been used for many years to determine binding sites of DNA-binding factors. To distinguish protein-specific interactions from various types of background, ChIP includes control samples consisting either of the chromatin input or the material resulting from non-specific binding of antibodies to chromatin. Many computational methods have been developed to identify ‘peaks’ from such data sets [13]. A previous study of peak finding methods developed for ChIP data has underscored the importance of the model describing the obtained data [14].

However, there are also differences. In contrast to ChIP, background samples are not always generated in CLIP experiments, largely because at the time when the field started it was unclear what an appropriate background should be. While at the DNA level, genes are generally represented in two copies per cell, the relative abundance of different RNAs in the cell varies over many orders of magnitude. Thus, abundant RNAs are likely to contaminate CLIP samples, leading to false positive sites, while binding sites in low-abundance RNAs may be completely missed. An approach to deal with this issue is to take advantage of crosslinking-induced mutations, identifying regions where such mutations have a higher than expected frequency [10,11,15,16]. This is not unproblematic, because mutations are introduced stochastically and the mutation-containing reads could also come from fragments crosslinked to proteins other than the one of interest in the experiment [17]. Another approach is to correct for the abundance of the RNAs based on RNA-seq data [18]. This is also not ideal, first because differences in sample preparation may lead to RNA-seq data not containing all the potential targets of the RBP, and second because the RNA-seq read coverage profile is not uniform, which will influence the quantification of the local background, and consequently the identification of CLIP sites. Finally, in the eCLIP variant of CLIP, the background coverage of transcripts by reads is inferred from a parallel sample that is prepared from the band corresponding roughly to the size of the protein of interest, obtained by omitting the immunoprecipitation step of sample preparation. This approach has the caveat that the size of the targeted protein varies from experiment to experiment, and so will the proteins that are contained in the isolated band. This makes it unclear whether the results obtained for different RBPs are of comparable accuracy.

As mentioned above, most RBPs bind their targets in a sequence-dependent manner, and sometimes in the context of specific structure elements [1,19]. For many RBPs, binding motifs have been inferred with both low and high-throughput approaches, and at least in some of these cases, there is good agreement between the RBP-binding motifs inferred from *in vitro* [20,21] and *in vivo* data [16,22]. However, in the most comprehensive database to date, ATtRACT [23], there typically are many motifs for an RBP, of widely variable information content and sometimes unrelated. A comprehensive database of RBP binding motifs determined from a consistent *in vivo* dataset, similar to those available for transcription factors [24,25], is still lacking.

A number of methods have been proposed for the identification of RBP binding peaks from CLIP data. Benchmarking of various subsets of these methods has revealed a few good performers, such as *CLIPper*, the tool developed for the analysis of above-mentioned eCLIP data, *omniCLIP* and *pureCLIP*, two recently published tools that use complex models to take advantage not only of the CLIP read coverage, but also of crosslinking-induced events (truncations or mutations) [15,18,22,26]. However, none of these tools provides an easy and robust end-to-end solution to the identification of binding sites and sequence motifs from CLIP data, and it has remained unclear how their accuracies compare.

To fill these gaps, we have developed RCRUNCH, a method that further aims to treat appropriately not only reads that map uniquely and contiguously to the genome, but also reads that map across splice junctions in mature mRNAs, as well as multi-mapping reads. The *de novo* motif discovery component of RCRUNCH, based on the well-established MotEvo tool [27], allows an immediate assessment of the quality of the results, including for the comparison of its genome/transcriptome or unique/multi-mapper based approaches. Using data for proteins with well-characterized sequence specificity, we demonstrate that RCRUNCH enables the reproducible extraction of binding sites, with higher enrichment in the expected motifs compared to the other tools. Application of RCRUNCH to the extensive eCLIP datasets generated in the ENCODE project [22], covering 149 RBPs, led to the construction of a comprehensive resource of *in vivo* binding sites and binding motifs of RBPs. RCRUNCH is available as an entirely automated tool from github (see Availability of Data and Materials).

## Results

### Automated CLIP data analysis with RCRUNCH

RCRUNCH is a workflow (Fig. 1a) for the automated and reproducible analysis of CLIP data, from reads to binding sites and motifs. It is written in the Snakemake language [28], observing the FAIR (findable, accessible, interoperable and reusable) principles [29]. The peak-calling module at the core of the workflow builds on the CRUNCH model [14] that was extensively validated on ChIP data. Along with the genome sequence and annotation files, the input to RCRUNCH consists of CLIP (foreground) sequencing reads, obtained from immunoprecipitation of a specific RBP, and background reads, which in most of the analyses reported here come from a size-matched input (SMI) control sample as in eCLIP. RCRUNCH’s default analysis mode uses reads that map uniquely to the genome, but multi-mappers and/or reads that map across splice junctions of mature mRNAs can also be included. For the latter case, RCRUNCH constructs a representative transcriptome composed of the isoform of each gene that has the highest abundance in the foreground sample. In a first step of its peak finding module, RCRUNCH identifies broad genomic regions whose coverage by reads is significantly higher in the foreground compared to the background sample (Fig. 1b, see Methods). A Gaussian mixture model is then applied to each of these regions to identify individual peaks and compute associated read enrichment scores (Fig. 1c). Peaks sorted by the significance of their read enrichment are then used in various analyses, including for the identification of enriched sequence motifs. The workflow provides extensive outputs such as the coordinates of the peaks, their enrichment scores and associated significance measure, enrichment values of known and *de novo* identified sequence motifs represented as positional weight matrices (PWM).

**Figure 1.**
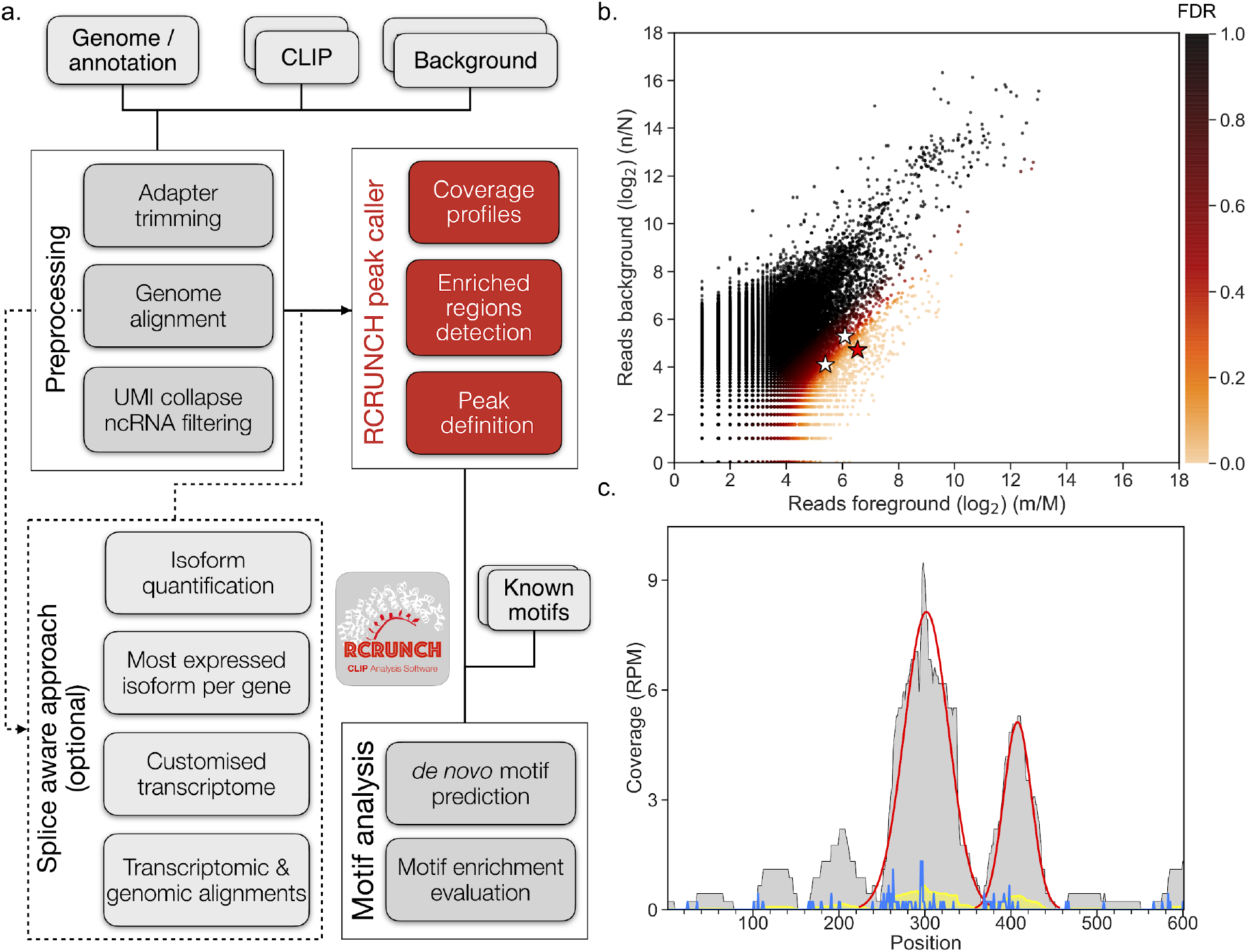
Schematic representation of RCRUNCH. **a**. Overview of the workflow. **b**. Scatterplot of the proportion of reads (log_2_) in individual genomic regions in a foreground (IP) sample generated by eCLIP for the PUM2 protein (replicate 2 of dataset ENCSR661ICQ from the ENCODE project, see Methods) and a corresponding background sample (SMI). Each dot corresponds to a 300 nucleotides-long genomic region. Marked in color are regions that are enriched in reads in the foreground relative to the background sample (FDR < 0.1, see color legend). Three genomic regions (zoom-ins in panel c) with various enrichment scores are marked with stars. **c**. Coverage of three overlapping genomic regions (highlighted in panel b and indicated at the top of the panel) by reads from the IP (light gray) and SMI (yellow) samples (RPM -reads per million). The significant peaks identified in the genomic region spanned by the three windows are shown in red. The blue line shows the number of read starts (5’ end of mate 2 reads) from the IP sample in the genomic region. Read starts indicate crosslinked nucleotides, where the reverse transcriptase falls off during sample preparation.

### Comparative evaluation of CLIP peak finding methods

By automating the analysis of CLIP data from reads to binding sites and motifs, RCRUNCH facilitates studies that rely on such data to a much larger extent than it was possible so far. To demonstrate its performance, we compared RCRUNCH with a few recently published and more broadly used tools for CLIP site identification. These were: *CLIPper*, the method used in the ENCODE project that generated the eCLIP data [22], already shown to supersede a few other methods [26], *PureClip* [18] and *omniCLIP* [15]. The latter two can use both the CLIP read density as well as the type and frequency of crosslinking-induced events in cDNAs to identify binding sites. As peak calling is not fully automated in *CLIPper*, for this tool we used the peaks provided by the ENCODE consortium [30,31]. The other tools were provided with genome-mapped reads obtained with the pre-processing module of RCRUNCH. As done in previous studies [15,18], in the comparative evaluation we used proteins for which the binding patterns and motifs have been extensively studied and are thus well understood [15,18,22]. These are hnRNPC, a splicing regulator that binds (U)_5_ sequences [32], PTBP1, another splicing regulator with a CU-rich binding motif [33], PUM2, a post-transcriptional regulator binding to UGUANAUA sequence elements [34,35] and RBFOX2, a splicing regulator known to recognize the sequence (U)GCAUG(U) [30]. For each of these proteins we applied RCRUNCH to the corresponding eCLIP samples in ENCODE, typically 2 replicates in each of two cell lines, and extracted the 1000 highest scoring sites from each sample (Supplementary Tables 1a-c).

The reproducibility of results obtained from replicate experiments is an important indicator of a method’s accuracy [12]. To estimate the agreement between sets of peaks, obtained from either replicate experiments or by different methods applied to the same dataset, we used the Jaccard similarity index (see Methods). The agreement of the top 1000 peaks inferred from two replicate experiments for the same RBP in a given cell line (Fig. 2a) was ∼20-40%, in the range reported for CLIP samples before [10]. RCRUNCH consistently provided values at the top of this range: for 5 of the 7 datasets RCRUNCH gave the highest agreement, and in the 2 cases when it did not, it was still a close second performer. Consistent with these numbers, most of the top 1000 sites from one replicate do not overlap with the top 1000 sites from the second replicate, though RCRUNCH consistently extracts a higher proportion of sites with high Jaccard index (Fig. 2b). We also asked how large is the overlap between the peaks reported by different methods. Although these numbers were generally lower than the overlap of peaks identified by one method from replicate experiments, RCRUNCH had the overall highest agreement with the other methods (Fig. 2c). The overlaps of RCRUNCH peaks from each sample with the peaks reported by the other methods for the corresponding protein are shown in Supplementary Table 1c. Finally, as the computational cost incurred by a tool is also an important factor in its adoption, we recorded the clock time for the peak calling step of all methods on the benchmarking data sets. For RCRUNCH the clock time was up to 3 hours (Fig. 2d), while *PureClip* needed up to 6 hours and omniCLIP up to 9 hours for an individual dataset/RBP.

**Figure 2.**
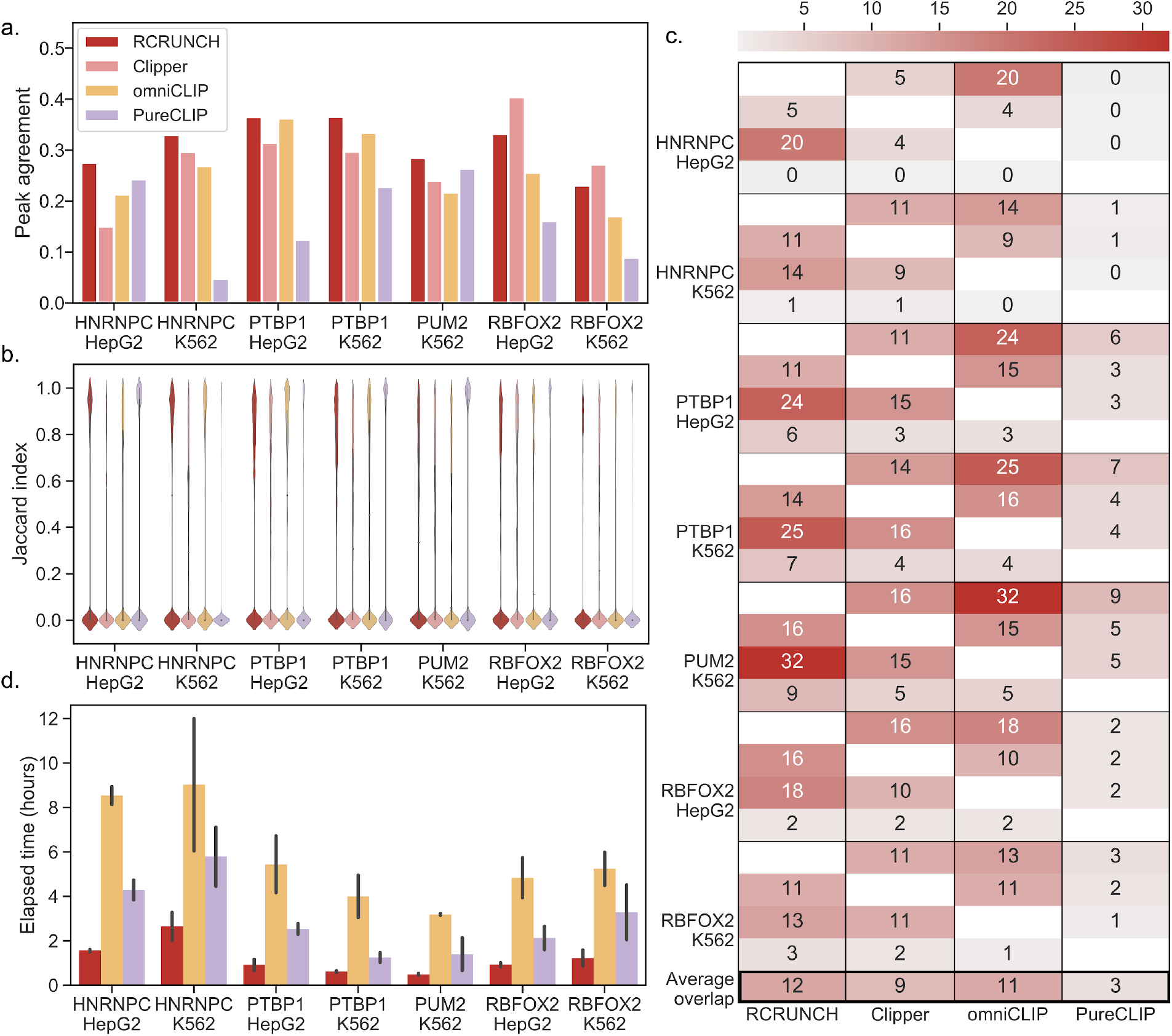
Comparison of CLIP peak calling methods. **a**. Barplot showing the Jaccard similarity index of the peaks identified by each computational method (shown in the legend) from replicate experiments. For all RBPs, except PUM2, data were available from two distinct cell lines, HepG2 and K562. **b**. Distribution of Jaccard indices of peak overlaps extracted by individual tools (color code as in panel a) from replicate samples. **c**. Heatmap showing the Jaccard similarity index of the peaks identified by two distinct methods for a given protein in a given cell line (in percentages, averaged over two replicate experiments). The methods are shown in order in the x-axis and the same order (RCRUNCH, CLIPper, omniCLIP, and PureCLIP) is used in each block (corresponding to one protein and cell line) on the y-axis. The average similarity of a method with any other method on all datasets is shown at the bottom. **d**. Barplot showing the running times of peak calling steps for RCRUNCH, omniCLIP, and PureCLIP. Error bars show the standard deviations from the 2 replicate runs.

Thus, RCRUNCH outperforms currently used methods both in terms of peak reproducibility between replicate experiments and in terms of running time. Moreover, RCRUNCH has, on average, the highest agreement with other peak finding methods, indicating that it capitalizes on some of the same information, while diminishing some of the biases of these other methods.

To further evaluate the quality of the detected peaks, we determined their enrichment in the motif known to be bound by the RBP targeted in each CLIP experiment. We extracted the literature-supported motifs for each RBP from the ATtRACT database [23] and calculated their enrichment in the 1000 top peaks (Supplementary Table 1) predicted by each method, relative to random genomic regions unlikely to be bound by the RBP (see Methods). As shown in Fig. 3a, the known motifs were indeed enriched in the peaks relative to background sequences, up to ∼5-fold, as observed before [22,36]. RCRUNCH peaks showed enrichment values at the high end of the achieved range (Fig. 3a) for all proteins/samples except hnRNPC. To verify that the peaks were indeed most enriched in the motifs known to be bound by the RBP (as opposed to any other motif), we further applied the Phylogibbs algorithm [37], to discover *de novo* the motif that is most overrepresented in the top peaks. Some of the *de novo* motifs were indeed similar to the expected ones, but they tended to be less polarized and more enriched (Fig. 3b-d). Strikingly, while Phylogibbs identified *de novo* motifs that were very strongly enriched in the hnRNPC peaks, these motifs did not have any resemblance to the expected (U)_5_ motif. To determine whether “truncated” reads, resulting from the reverse transcriptase falling off at the crosslinked nucleotide, allow a more accurate identification of RBP-binding motifs than the peaks in read coverage, we implemented the “RCRUNCH crosslink” variant, in which RBP binding sites are extracted from around the most crosslinked position within each peak (position where most reads start), in contrast to the “RCRUNCH peak center” discussed so far, in which sites are extracted relative to coverage peak centers. RCRUNCH crosslink clearly recovered the hnRNPC-specific (U)_5_ motif (Fig. 4d) and further improved the identification of the RBFOX2-specific UGCAUG motif, while the recovery of the PUM2 and PTBP1-specific motifs was unaffected.

**Figure 3.**
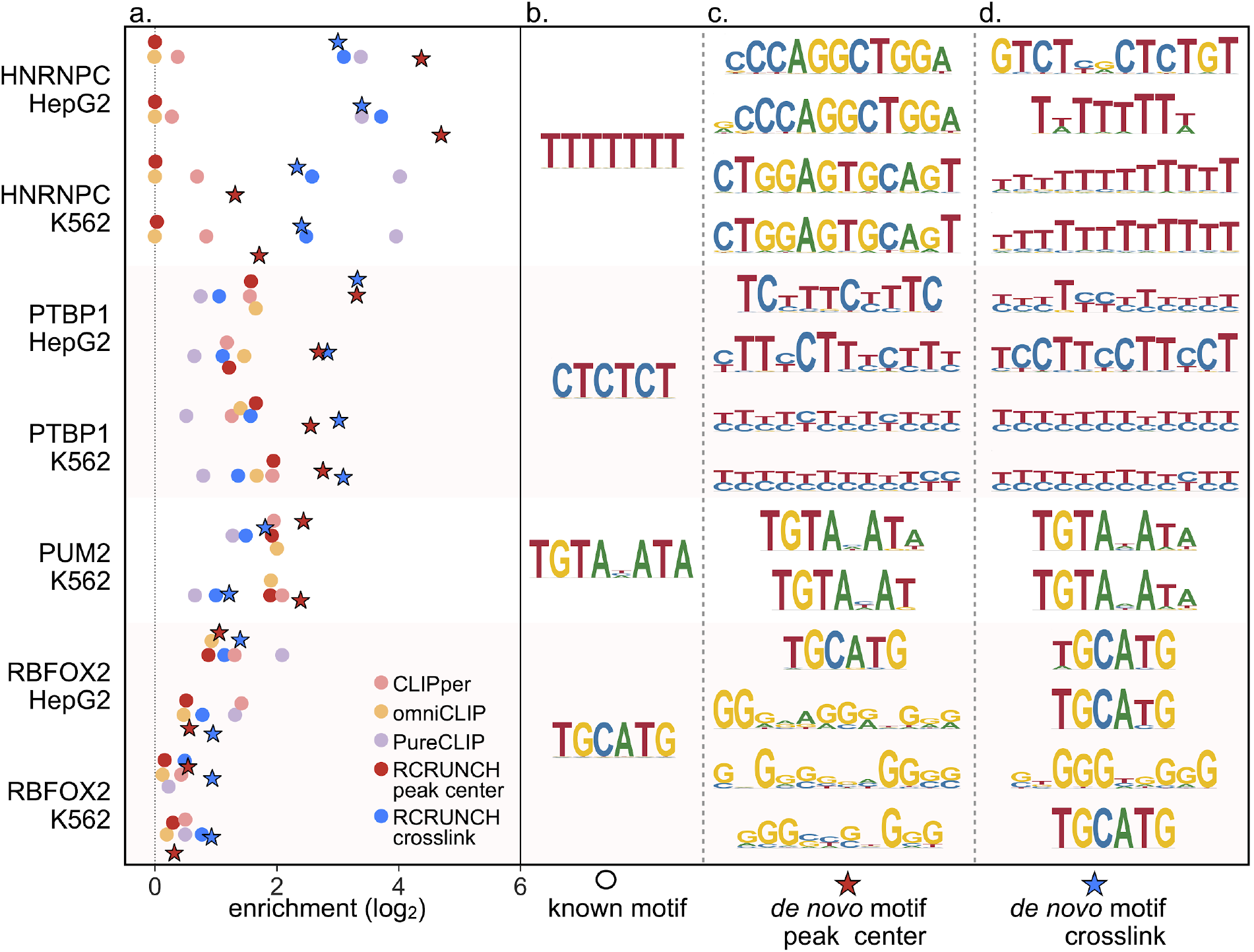
Enrichment of known and *de novo* sequence motifs in the eCLIP data of individual RBPs. **a**. Enrichment scores computed by comparing the frequency of known motifs among the top CLIP peaks identified by the indicated method with the frequency in background regions (random subsets of regions that were least enriched in CLIP reads, see Methods). Each peak finder is shown in a different color. The enrichments of the *de novo* motifs are indicated by stars, red for RCRUNCH peak center and blue for RCRUNCH crosslink. **b**. The known RBP-specific motifs from ATtRACT [23] that were used in the analysis. **c**. Most significantly enriched *de novo* motif predicted by Phylogibbs [37] in the “RCRUNCH peak center” sites from each sample. **d**. Most significantly enriched *de novo* motif predicted by Phylogibbs in the “RCRUNCH crosslink” sites from each sample.

**Figure 4.**
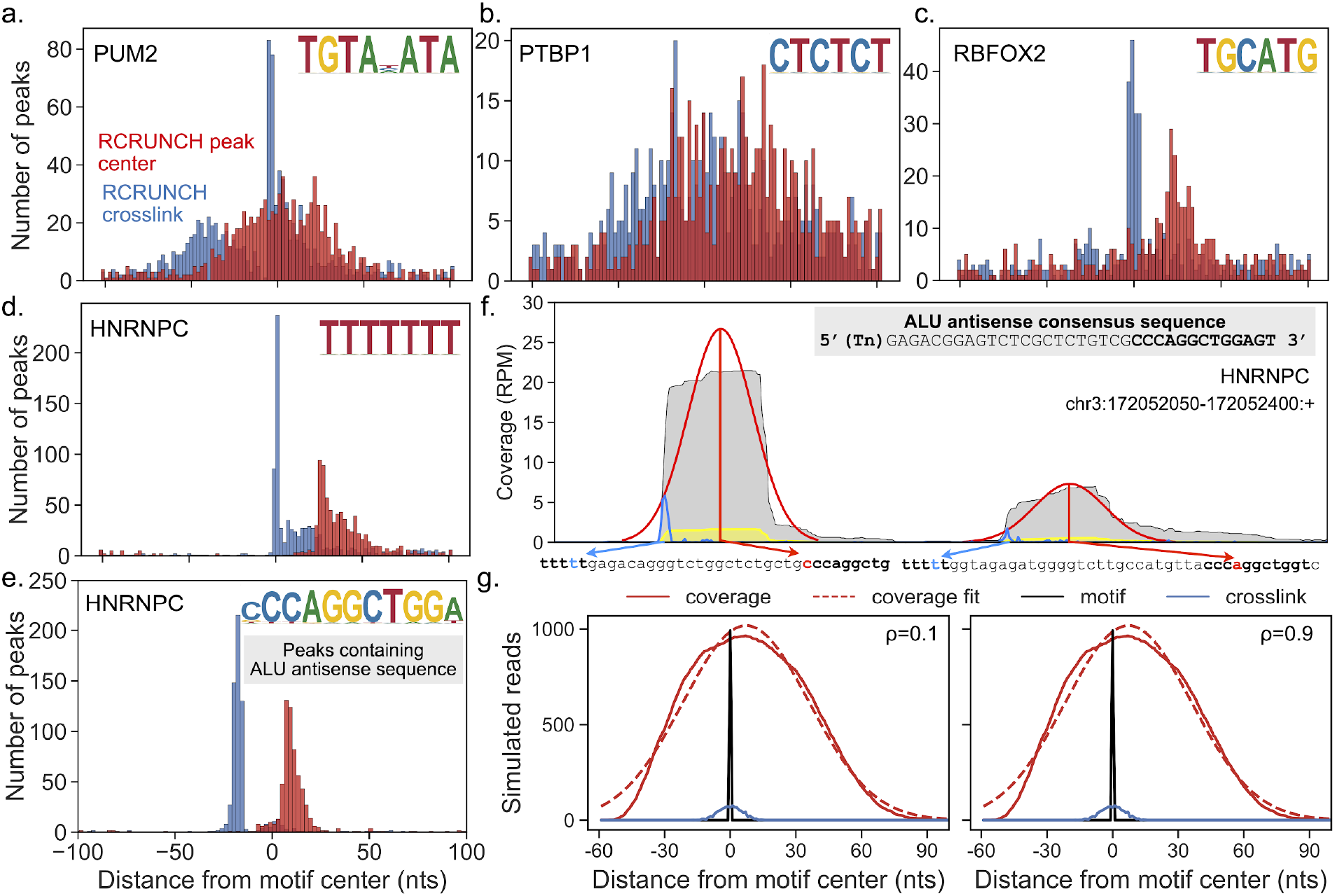
Different configuration of binding and crosslinking across RBPs. **a-d**. Histograms of distances between the cognate motifs of RBPs and the centers of the read coverage peaks (RCRUNCH peak center, in red) or between the motifs and the most frequent read start position in a given peak (RCRUNCH crosslink, in blue). The top 1000 peaks (in the order of their z-score) from one of the available samples for PUM2 (a), PTBP1 (b), RBFOX2 (c) and hnRNPC (d) were extracted. The cognate motif with the highest posterior probability given the RBP’s PWM was determined and the peak was retained only when the posterior was at least 0.3. **e**. For hnRNPC we carried out the same analysis relative to the Alu-related motif. **f**. Example of two hnRNPC binding peaks located on Alu antisense elements. The library-size-normalized read coverage in the eCLIP IP sample is shown in gray, while the coverage in the corresponding SMI sample is shown in yellow. The fitted Gaussian peaks predicted by RCRUNCH are shown in red, while the distribution of reads starts in this region is depicted with the blue line. The most frequent read start within a coverage peak is chosen by RCRUNCH crosslink as the crosslink position and the corresponding nucleotide is indicated here by the blue arrow. The red arrow links the peak center to the corresponding nucleotide within the Alu antisense element. RPM -reads per million. **g**. Results of CLIP experiment simulations, showing the read coverage profile (full red line), corresponding Gaussian fit (dashed red line), and frequency of crosslinks (blue) with respect to the RBP-specific motif (black, centered on position 0), for low (0.1, left) and high (0.9, right) probability of reverse transcriptase readthrough (*ρ*).

These results demonstrate that the peaks predicted by different methods are enriched to fairly similar extents in the expected motifs, though RCRUNCH has the most reliable high enrichments. Furthermore, the *de novo* motifs identified by RCRUNCH are more enriched in the peaks than the known motifs, even when they appear quite similar. For some proteins, specifically hnRNPC and RBFOX2, the read starts enable a more precise identification of RBP-specific binding motifs, while for others, like PTBP1 and PUM2, the coverage peak centers are equally informative.

### RCRUNCH helps elucidate how RBPs interact with and crosslink to RNAs

To better understand why the sites extracted from around coverage peak centers contain the RBP-binding motif for some proteins but not for others, we carried out the following analysis. Within each of the 1000 most significant peaks based on the enrichment in reads we identified both the location of the highest-scoring match to the RBP-specific motif and the crosslink position, where most reads started (Supplementary Table 1). Then, we anchored the peaks on the center of the motif match (position 0), and constructed the histograms of distances between the crosslinks and motifs, and between coverage peak centers and motifs. As already suggested by the observations from the previous section, the relationship between these two histograms is highly dependent on the studied RBP (Fig. 4a-d). For PUM2, RBFOX2, and hnRNPC, crosslink positions strongly co-localize with the RBP-binding motif, while for PTBP1 this is not the case. In contrast, the peak centers show weak co-localization with the binding motif of PUM2 and PTBP1, but occur clearly downstream of the binding motifs for RBFOX2 and hnRNPC.

In the case of hnRNPC, highly enriched motifs were recovered around peak centers as well, and these motifs were very different from the expected (U)_5_ (Fig. 3c). A literature search revealed that these motifs correspond to the Alu antisense element (AAE) [38], consistent with the reported function of hnRNPC in suppressing the exonization of these repetitive elements [39]. Computing the peak center - motif and crosslink - motif histograms relative to the AAE showed that hnRNPC binding sites containing AAEs have a very specific configuration, crosslinking occurring upstream of the AAE, within the U-rich motif, leading to CLIP read starts in this region, while the peak in read coverage is on the AAE (Fig. 4e,f). To better understand how these patterns relate to the specific interaction of an RBP with RNAs, we carried out a simulation of an RBP binding to its cognate motif, crosslinking to the RNA and protecting an extended region of the target from RNase digestion. As the efficiency of crosslinking depends on the identity of both the nucleotides and the amino acids that participate in the RNA-RBP interaction [40], we simulated three scenarios, corresponding to the nucleotide with the highest efficiency of crosslinking being located upstream, within or downstream of the RBP-specific binding motif. For each of these cases we simulated scenarios where the probability *ρ* of reverse transcriptase reading through the crosslinked nucleotide is very low, intermediate or very high (Fig. S1). We found that when the readthrough probability is low, the crosslink is a better indicator of the binding motif than the peak center (Fig. 4g). In contrast, when the readthrough probability is moderate to high, the crosslink position and the coverage peak center are located at comparable distances from the RBP-binding motif, so that either could be used to identify the RBP binding site (Fig. 4g). Thus, the motif-crosslink and motif-peak center distance relationships that we observed for the selected proteins indicate that the probability that the reverse transcriptase reads through the RBP-RNA crosslink is much lower in the case of RBFOX2 and hnRNPC compared PUM2 and PTBP1, leading to a more precise identification of the binding motif when binding sites are centered on the crosslink position. That “truncations” are not always good indicators of the location of the binding motif has been reported before, a main reason being the variation in fragment size between experiments [41,42]. This is unlikely to be the cause in the variation we observe here because the fragments sizes were very similar (∼10 nucleotides, Suppl. Table 2) between samples.

These results suggest that model-driven analyses of CLIP data, taking into account the architecture of protein-RNA interactions, could further improve the identification of binding sites and the interpretation of the observed binding patterns. Furthermore, the variant RCRUNCH workflows provide a flexible platform to explore the architecture of RBP-RNA interaction sites.

### RCRUNCH variants enable detection of specific classes of RBP targets

Tools for CLIP data analysis focus almost exclusively on reads that map uniquely to the genome, leaving out multi-mapping reads or reads that map across splice junctions, which are more challenging to map and quantify correctly. To provide users with the opportunity to investigate RBPs that specifically bind to repetitive elements or mature mRNAs, we have implemented and evaluated a few variations of the RCRUNCH workflow. Specifically, we have implemented the option of identifying binding sites that are located in the immediate vicinity of exon-exon junctions in mature mRNAs, as well as the option of using reads that map to multiple genomic locations (multi-mappers). In the first situation, some reads end up mapping to the genome in a split manner, partly to the 5’ exon and partly to the 3’ exon. This in turn can lead to multiple distant peaks, with the RBP-binding motif being present at only one or perhaps neither of these peaks. Finally, the question of an appropriate “background” for estimating the enrichment of reads in CLIP samples is still open [12]. Aside from the SMI control used here, the relative abundance of mRNAs (estimated based on RNA-seq data) is sometimes taken into account [43]. RCRUNCH allows an easy incorporation of different types of background, provided as an appropriate file with sequenced reads. Here we used the mRNA-seq data generated for the specific cell lines included in the ENCODE project. We benchmarked the performance of RCRUNCH variants on the proteins chosen at the beginning of our study. In all cases we extracted sites anchored at the crosslink position within each peak and compared the “standard” RCRUNCH crosslink with individual variants across a few different measures. These measures were: the number of significant sites (at FDR = 0.1) identified in a sample (Fig. 5a-c), the enrichment of the known motif in the top 1000 sites identified in each sample (Fig. 5d-f), and the similarity of the known motif of the RBP to the de novo motif identified from the top 1000 peaks of a given sample (Fig. 5g-i). While for the RCRUNCH transcriptomic and multi-mapper approaches the results were very comparable with those of the standard RCRUNCH, the choice of RNA-seq as background results in a strong decrease in performance. Including multi-mappers or splice junction reads led to the recovery of somewhat fewer sites, but the quality of the peaks, measured in terms of their enrichment in motifs, was not affected.

**Figure 5.**
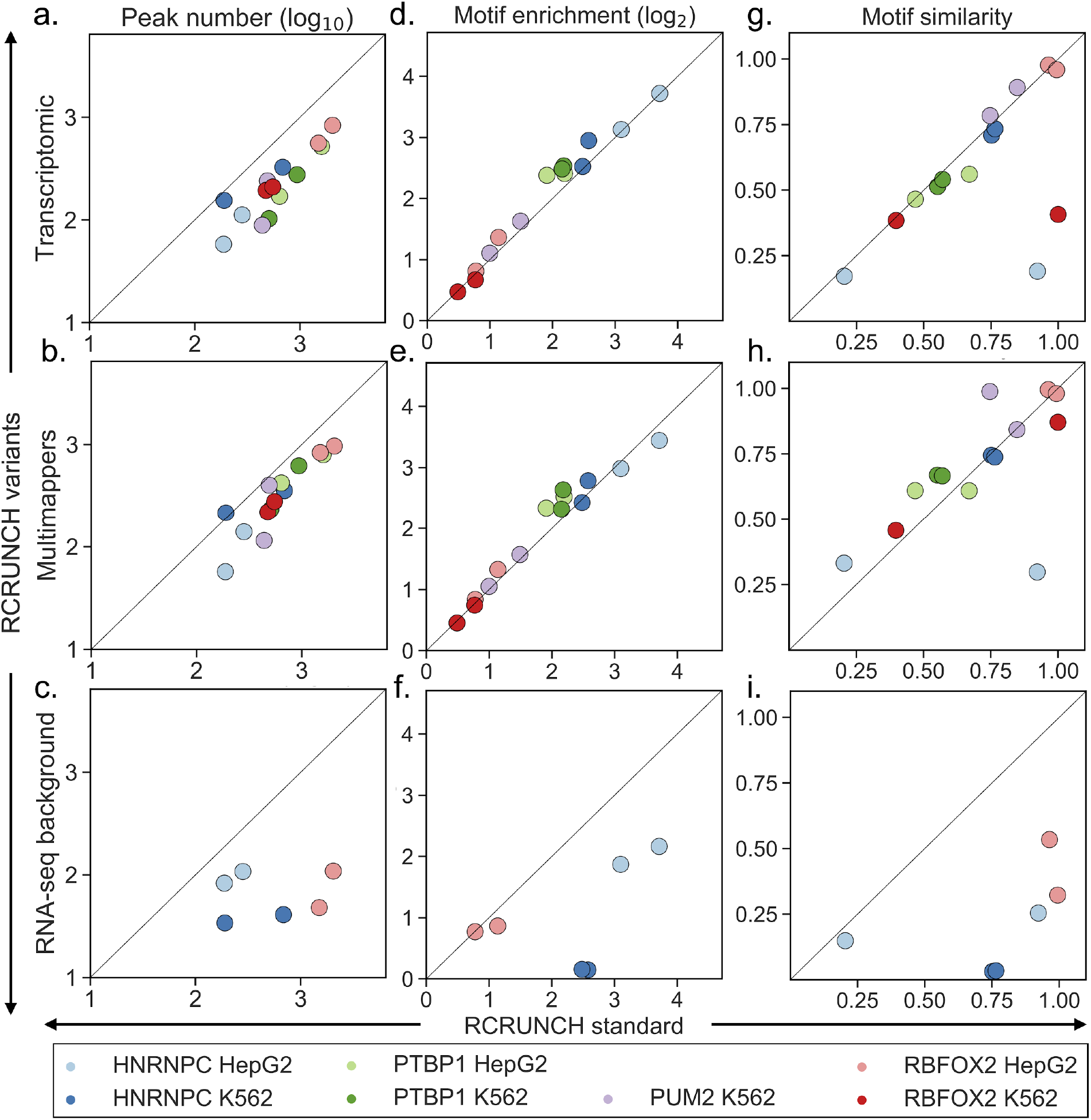
Performance evaluation of RCRUNCH variants. “RCRUNCH standard” refers to peaks identified based on reads that map uniquely to the genome, and binding sites that are always centered on the most frequent crosslink position in each peak. Rows correspond to variants of the RCRUNCH workflow, while columns show different metrics used to compare each of the variants with the “standard” RCRUNCH: **a-c**. Number of significant peaks in a sample (at FDR < 0.1); **d-f**. Enrichment of the known motif of the RBP that was assayed in a particular sample; **g-i**. Similarity of the motif identified *de novo* from the top peaks of a given sample and the known motif for the respective RBP. In the case of the RNA-seq background variant, some samples did not yield any significant peaks (FDR = 0.1). These samples are therefore not represented in the plots.

These results demonstrate that RCRUNCH is a flexible and performant method for CLIP data analysis. The choice of background is important, and in the case of eCLIP, the SMI control samples provide a more appropriate background for estimating read enrichment in binding sites than the RNA-seq samples.

### A compendium of RBP binding motifs inferred from eCLIP data

Although various analyses of the ENCODE eCLIP datasets have been carried out, a consolidated compendium of binding motifs inferred for individual proteins from these data is not available. To fill this gap, we have applied RCRUNCH (both peak center and crosslink variants) to all available samples in these dataset, determined peaks that are enriched in reads from IP relative to the SMI, inferred the most significantly enriched sequence motifs and finally, for each RBP, identified the motif with the highest average enrichment across all samples corresponding to the RBP (Supplementary Table 3). The distribution of the number of binding sites per RBP, as shown in Fig. 6a, indicates that two thirds of the samples yielded more than 100 binding sites, with few samples (for the HNRNPL, AGGF1, DDX3, TARDBP proteins) yielding thousands of sites. As may have been expected, a known binding motif in ATtRACT is indicative of the protein having a high number of binding sites (Fig. 6a-b). We next calculated the average Jaccard similarity index of peaks identified from pairs of samples, either corresponding to the same protein, or to different proteins (Fig. S2). We also carried out an analysis of the motifs enriched in samples for an individual RBP (Fig. S3), ultimately identifying the motif that best explains the entire data obtained for a given protein (highest sum of log-likelihood ratios across all samples, Fig. 6c-d). Of 149 proteins, 86 yielded an enriched motif in our analysis, and 26 of these already had a specific motif in the ATtRACT database. 21 of the proteins for which we could not identify an enriched motif in this study were covered by the ATtRACT database (Fig. 6b). The heatmap of peak overlaps shows good consistency among different experiments involving the same protein (Fig. 6c, diagonal), and also highlights interesting cases of proteins that bind to similar regions, in many cases because the proteins take part in the same multi-molecular complex. For example, we found high overlaps between the sites of splicing factors U2AF1 and U2AF2 [44,45], of the DXH30 and FASTKD2 proteins involved in the ribosomes biogenesis in the mitochondria (Fig. S2) [46], of the DGCR8 and DROSHA components of the miRNA biogenesis complex [47] and a few others (Fig. 6c). These results lend further support to the notion that our method recovers expected signals in the eCLIP data. Additional per sample analyses are shown in Fig. S2. In brief, we identified enriched motifs in 90% of the samples, similar between the RCRUNCH crosslink and RCRUNCH peak center approaches (Fig. S4a), but in many cases the enrichments were small. As we have seen for the benchmarked proteins, the enrichment of the *de novo* identified motif was higher than the enrichment of the known motif for the studied protein (Fig. S4b). We further calculated the similarity (see Methods) between the known and *de novo* motifs, the latter obtained either from RCRUNCH crosslink or RCRUNCH peak center-predicted binding sites. We found that around 80% of the samples yielded motifs with at least 0.4 similarity to the known motif, which is a much higher proportion than when comparing random pairs of motifs. The similarity was slightly higher when sites were identified by RCRUNCH crosslink (Fig. S4c). Given the large number of motifs identified for RBPs that are not represented in ATtRACT, we asked whether these motifs are reproduced between replicate samples of an RBP. Indeed, the similarity of motifs obtained from replicate samples was similar for proteins with and without a known motif, and it correlated with the number of sites inferred from the samples (Fig. S4d). De novo motifs that were reproduced between replicate samples were also generally similar to known motifs for the corresponding protein (Fig. S4e).

**Figure 6.**
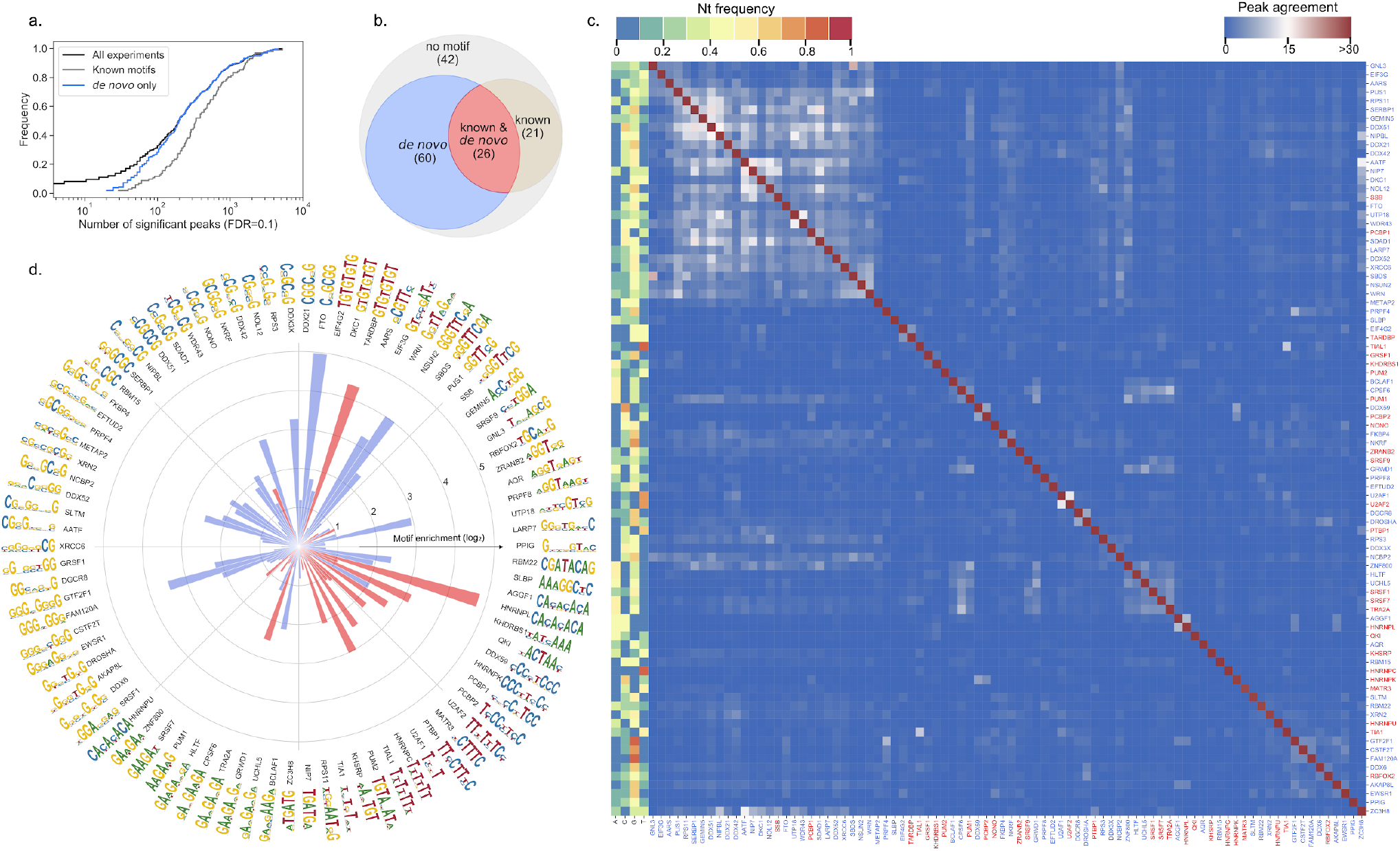
RCRUNCH results for all ENCODE eCLIP data currently available. **a**. Cumulative distribution of the number of significant binding sites detected per experiment (FDR threshold=0.1, in black). Cumulative distributions are also shown separately for samples corresponding to proteins with a known binding motif (gray) and to proteins for which no known motif is available in ATtRACT, but one was found by RCRUNCH (blue). **b**. Venn diagram summarizing the motif inference in the identified peaks. We distinguished four categories of proteins: for which (1) no motif is known and also no enriched motif was identified in this study (grey), (2) a *de novo* motif was found for a protein for which no motif is given in ATtRACT (blue) (3) a *de novo* motif was found for a protein with a known motif in ATtRACT (coral) and (4) a motif is known, but none was identified *de novo* (sand color). **c**. Heatmap of mean peak agreement across RBPs. Only RBPs for which an enriched motif (see Methods) was found are included. The agreement is calculated as the Jaccard index of the nucleotides (nts) in the peaks, where the intersection of two sets of peaks is the number of nts covered in both sets, while the union is the number of nts covered in at least one of the two sets. The color range is capped at a similarity of 0.3 to make the clusters more easily distinguishable. The top peaks are taken according to the FDR threshold (0.1), extending by 20 nts upstream and downstream from the crosslink site. Since there are multiple replicates per RBP, the mean of pairwise Jaccard indices over all combinations of sample pairs are used here. The colors on the left indicate the relative frequency of each nucleotide type averaged over all positions of the PWM. **d**. Polar projection of the enrichment of *de novo* motifs inferred for individual RBPs from all peaks with FDR >0.1 extracted from the ENCODE samples. Only RBPs for which an enriched motif (see Methods) was found are included. The color of the bars indicate whether the respective RBP already has a known motif in ATtRACT (coral) or not (blue).Supplementary data

These data indicate that the motifs identified from RCRUNCH-extracted CLIP peaks are reliable, conform with prior knowledge, and explain the binding data better than the motifs that are currently available in databases. Altogether, the compendium that we have constructed (Supplementary Table 3, see also Availability of Data and Materials), provides sequence specificity data for 86 RBPs, thus being, to our knowledge, the most extensive collection of *in vivo*, reproducibly-identified, consensus RBP binding motifs.

## Discussion

Within over a decade of development and application, CLIP has provided a wealth of insight into the RNA-binding protein-dependent regulation of cellular processes such as RNA maturation, turnover, localization, and translation. The large collection of RBP-centric high-throughput datasets that has been generated as part of the ENCODE project [48] is broadly used to unravel the functions of RBPs, many of which were only recently found to bind RNAs. As other types of high-throughput data, CLIP also requires dedicated computational analysis methods. In this context, RCRUNCH makes the following main contributions.

First, it is the first completely automated solution to CLIP data analysis, from reads to binding sites and sequence motifs, with accuracy and running times that compare favorably to those of the most broadly used tools to date. Key to this performance are the enrichment-based prioritization of genomic regions likely to contain RBP binding sites, and the model for evaluating this enrichment in individual sites. These are based on the CRUNCH tool for analyzing ChIP data, and we find that they largely apply to CLIP data as well (Fig. S5).

Second, to accommodate the variability of target types across RBPs, RCRUNCH goes beyond the typical approach of using uniquely genome-mapped reads, allowing the inclusion of multi-mappers and/or of reads that map across splice junctions. Both of these situations make it difficult to determine the locus of origin of the reads and may lead to a decreased accuracy of binding site inference. Nevertheless, for RBPs that specifically bind repeat elements, or in the vicinity of splice junctions, taking into account such reads and appropriately defining the peaks in read coverage is a must. The variant RCRUNCH workflows fulfill this need. RCRUNCH multi-mapper considers reads that map to a maximum number of genomic locations (specified by the user), distributing the reads equally among the loci with maximum alignment score. Compared to the run that only allowed uniquely mapped reads, this approach yields binding sites with similar enrichments in the expected sequence motifs of the benchmarked proteins, including those with repetitive binding motifs like PTBP1. More sophisticated models, e.g. applying an expectation/maximization approach to read-to-locus assignment, have been proposed before [49], but they come at the cost of increased running times, and have not been broadly adopted. The RCRUNCH transcriptome variant was designed to address another special case, namely that of RBPs that bind predominantly mature mRNAs. Many RBPs are located in the cytoplasm where they orchestrate RNA traffic and localization [36]. As internal exons of human transcripts are relatively short, ∼147 nucleotides [50], it is likely that CLIP reads for RBPs that bind to these exons will cover exon-exon junctions [51]. Recent generation alignment programs such as STAR [52] can carry out spliced alignment. However, reads that span splice junctions will give rise to multiple peaks, in the 5’ and the 3’ exons that flank the splice junction. This will affect the accuracy of binding site and motif identification. A way to circumvent the issue is to map the reads to transcripts instead of genome sequences. This is not without drawbacks. First, given that it is not generally known a priori whether the RBP of interest binds mature RNAs or other classes of transcripts, a hybrid strategy will need to be adopted, to allow the identification of binding sites in introns as well as in mature mRNAs. Second, given the large number of possible isoforms per gene, accurate assignment of reads to isoforms and peak identification in multiple isoforms are not trivial. In RCRUNCH transcriptome we have implemented a general hybrid strategy to capture binding sites across all types of types of transcripts, including pre-mRNAs and mature mRNAs, without incurring large computational costs. Namely, taking advantage of the observation that individual cell types express predominantly one isoform from a given gene [53], we first determine which of the known isoforms of each gene has the highest expression level in the IP sample. We then use these isoforms as pseudo-chromosomes, assigning reads to the best-scoring loci, but with priority given to the spliced isoforms over genomic loci. This approach gave good results for the benchmarked proteins, including PUM2, which is known to bind to mature mRNAs [54], but overall, the number of sites that spanned splice junctions was small for the benchmarked proteins. On the other hand, for splicing factors we identified many sites in the vicinity of splice sites, as expected, indicating that these data can be studied further to determine to what extent these splicing factors remain associated with the mature RNAs (Fig. S6). Nevertheless, our exploration of ways to handle reads that originate in various categories of targets was by no means exhaustive and this could be a direction for further development of the workflow.

Third, our analysis of the ENCODE eCLIP data yielded enriched sequence motifs for 86 RBPs. These were selected using a uniform procedure, based on the maximum enrichment across all samples available for a given RBP, in contrast to resources such as ATtRACT, which contain multiple, heterogeneous RBP-specific sequence motifs obtained with a wide range of techniques. While the eCLIP datasets were the focus of various previous studies (e.g. [16,22,36]), a compendium of reproducible binding motifs inferred from these data sets is not available. Moreover, although RBPs typically bind to a defined sequence (or sometimes structure) motif, the binding specificity of RBPs inferred from eCLIP data has been described in terms of collections of short motifs [16,36], for reasons that remain unclear. The main aim of the work presented here was to provide a uniform procedure for inferring RBP sequence specificity from binding data, and a resource of RBP-specific motifs similar to those available for transcription factors (e.g. [55]). For RBPs that have been extensively studied, the motifs that we identified *de novo* from the CLIP peaks conform with prior knowledge, though they differ in quantitative detail. Moreover, the *de novo* motifs have higher enrichment in the peaks compared to the known motifs, which may indicate context-specific contributions to the binding affinity. Overall, we identified enriched sequence motifs for 86 proteins, 60 of which are not represented in the ATtRACT database. In some cases, the most enriched motif in a given sample was not the one known to be bound by the corresponding protein, as observed before [36]. Repetitive motifs (G/G&C/C-rich) were occasionally found to be enriched in various samples, and this enrichment was also reproduced in replicate samples for the same protein. This raises the question of whether these motifs represent some sort of non-specific background in IP samples [18]. However, we did not find a larger overlap among the binding sites containing such motifs relative to binding sites of randomly chosen pairs of proteins (Fig. 6c). In fact, overlaying the pairwise overlap data with data on protein complex composition revealed compelling cases of high overlap for proteins of the same complex such as the spliceosome, the pre-rRNA processing complex, a paraspeckle-related complex and others (Fig. S2). Thus, our analysis does not support the concept that general non-specific background in eCLIP leads to similar motifs for unrelated RBPs, though it will be interesting to investigate further the functional significance of the identified motifs.

Finally, our analysis of the motif-crosslink and motif-peak center distances revealed distinct RBP-dependent patterns. Most striking was the peak in coverage observed over the AAEs in the hnRNPC eCLIP. HnRNPC binds (U)_5_ elements [9], while the read starts, indicative of crosslink positions, were located in a U-rich region upstream of the AAE. Our simulation of a CLIP experiment suggests that the hnRNPC data is quite unusual relative to data for other RBPs. HnRNPC has a large number of binding sites in Alu antisense elements that have a conserved consensus extending much beyond the U-rich element. Motif finding methods will identify this consensus as extremely enriched, more so than the much shorter (U)_5_ motif. The strong colocalization of the most frequent crosslink position within a peak and the RBP-specific motif supports the notion that the reverse transcriptase has a high propensity to stop extending the cDNA when it encounters the RNA-RBP crosslink [56]. However, our analysis also suggests that the readthough probability varies substantially between RBPs, being very low for hnRNPC, and relatively high for other proteins like PTBP1 and PUM2. This highlights the importance of a flexible but principled approach to binding site and motif identification, using a general measure of performance such as the motif enrichment score. This is what RCRUNCH implements, running in parallel the standard and crosslink approach and reporting the motif enrichments to allow the user to determine the approach that yields the most consistent results. This is because the configuration of RBP-RNA interactions varies across RBPs, influencing the nature of the reads that are captured in the CLIP experiment. Although beyond the scope of our present work, exploration of the ENCODE data set with a simulation-driven approach may yield further insight into the interactions of individual RBPs with their binding sites.

To ensure comparability of various tools, we used the metric provided by each of the tools to select the same number of top sites for further analyses. The question may arise whether the number of selected sites influences our conclusions. We did not find this to be the case (Fig. S7). Specifically, the relative ranking of motif densities reported by different methods largely held for a number of top sites between 0 and 1000.

The ENCODE project used the eCLIP approach, and thus eCLIP datasets are by far the most extensive. Aiming to obtain a comprehensive catalog of sequence motifs, we have therefore focused our analysis on these datasets. However, RCRUNCH is not specifically tailored to eCLIP. For example, it can also use single-end sequencing data, with or without UMIs, generated by eCLIP or another protocol. What is important though is that background samples, not enriched for the protein of interest, are available. To illustrate this, we have analyzed data generated by PAR-CLIP for the fly CNBP protein (Fig. S8), obtaining the expected sequence motif. In another case, PAR-CLIP data for the PUM2 protein, where we tried to use as background RNA-seq data from similar experimental systems, the read counts per window did not follow the expected distribution, and we were not able to compute meaningful enriched regions (Fig. S8).

## Conclusions

Our study provides a general, end-to-end solution for CLIP data analysis, starting from sequenced reads and ending in binding sites and RBP-specific sequence motifs. The tool compares favorably with the most broadly used tools to date, and further extends the type of reads that can be analyzed, to multi-mapping and split-mapping reads. By applying RCRUNCH to the entire ENCODE set of samples available to date, we provide a compendium of reproducibly enriched sequenced motifs for 86 RBPs, of which only 26 are represented in extensive databases available today, such as ATtRACT. Finally, our simulations suggest that the architecture of RBP-RNA interactions imposes strong variation in the probability of identifying the precise position of crosslinking from CLIP data.

## Methods

### Inputs to RCRUNCH

RCRUNCH performs its analysis on at least one paired-end or single-end, stranded CLIP sample, and a corresponding background sample (which could be the SMI from eCLIP experiments or RNA-seq) both provided in fastq format. All necessary parameters for the run such as sample file names, adapters, fragment size, presence of UMIs etc. should be provided in a config file, a template of which can be found in the repository along with a test case. The tool also requires the genome sequence fasta file and Ensembl [57] gtf annotation of the corresponding organism, also given in the config file. RCRUNCH can additionally perform some optional analyses, one being the filtering out of sequences that correspond to specific non-coding RNA biotypes and the other being the enrichment analysis for known sequence motifs. If these options are chosen, paths to corresponding files should be provided in the config file, according to the instructions in the README file accompanying the software.

### Read preprocessing

#### Adapter removal

3’ and 5’ adapters for read1 and/or read2 specified in the config file are trimmed with Cutadapt [58].

### Alignment of reads to reference genome

The alignment of reads to the reference genome is done with STAR [52], disabling the soft-clipping option. Some of the options to STAR differ from the standard value, to allow the alignment of short reads with only few mismatches (outFilterScoreMinOverLread 0.2, --outFilterMatchNminOverLread 0.2, outFilterMismatchNoverLmax 0.1). Multi-mapper reads (that map equally well - same number of errors - to multiple regions in the genome) can be included in the analysis by setting the ‘multimappers’ field in the config file to the desired number of equivalent mappings to consider for a read. In this case, reads that map to at most ‘multimappers’ locations in the genome are counted towards each of these locations with a weight of 1/’multimappers’.

### Removal of reads from abundant non-coding RNAs

Reads derived from some non-coding RNAs (e.g ribosomal (rRNAs), transfer (tRNAs) and small nuclear RNAs (snRNAs)) are abundant in many CLIP samples and thus believed to be largely contaminants [22]. Frequently, these abundant RNAs are also encoded in highly repetitive genomic loci. For these reasons RCRUNCH allows the option of selective removal of reads mapping to ncRNAs, based on the annotation from RNAcentral [59]. For this, the user will need to provide a gff3-formatted file for the appropriate species, which can be downloaded from RNAcentral. Specific biotypes of ncRNAs can be selectively removed by filling out the ‘ncRNA_biotypes’ option in the config. The names of reads that overlap in the genome with any of the selected ncRNAs specified by the user, are saved in a list. This is then used as input in the FilterSamReads function of the Picard software [60] to remove the reads from the alignment file that passed to downstream analysis.

### Removal of PCR duplicates

PCR amplification is a well-established source of error in the estimation of transcript counts [61]. However, different CLIP protocols differ in whether and how they deal with this issue. Accordingly, RCRUNCH offers multiple options. The default is to not carry out any PCR duplicate removal, which can be specified by choosing ‘standard’ as the value for the ‘dup_type’ fields in the config file. Alternatively, RCRUNCH can take advantage of Unique Molecular Identifiers (UMIs), which are introduced by ligation of a DNA adapter containing a random oligonucleotide (the UMI or randomer) to the cDNA fragments, as done in eCLIP [22]. As the UMI is preserved during PCR amplification, it can be used to identify reads that are copies of the same initial fragment. To remove PCR duplicates we use UMI-tools [62], which assumes that the UMI sequences are suffices of the read names. However, data from the ENCODE project has the UMIs as prefixes to the read names. Thus, we use a specific rule to make this transformation, which is controlled by the field ‘format’ in the config file, and can be either ‘encode’ or ‘standard’. If standard is chosen, no reformatting occurs and it is up to the user to make sure the format of the fastq files they provide is compatible with UMI-tools processing. Finally, if the sample preparation did not include the addition of UMIs, RCRUNCH can still attempt this removal via the deduplication function of STAR [52] via filling out the ‘dup_type’ option with ‘duplicates’. If no duplicate removal is desired then the ‘dup_type’ can take the option ‘with_duplicates’.

### Additional preprocessing steps for the ‘RCRUNCH transcriptome’ approach

If the user chooses the transcriptomic mode of RCRUNCH (‘method_types’ as ‘TR’ in the config), a few additional steps are needed to identify reads that map across splice junctions. First, reads are aligned to the genome (as described above), and the alignments are used to remove PCR duplicates and possibly ncRNAs. The remaining read alignments for the foreground sample are used by the Salmon software [52,60,63] to select the most expressed transcript isoform for each gene and construct a dataset-specific transcriptome. The reads that were selected in the first step are aligned to the transcriptome, after which the genome and transcriptome alignment files are jointly analyzed to identify the highest scoring alignment (AS score in the bamfiles) for each read. If the AS score for the transcriptome alignment is greater than the AS score of the genome alignment -3, the alignment to the transcriptome is selected. We chose this criterion rather than requiring the transcriptome alignment score to be strictly better than the genome alignment score to conservatively assign the reads preferentially to the transcriptome. Peaks are then detected either on the genome or the transcriptome, treating individual transcripts as we treat chromosomes. This approach allows us to detect and properly quantify RBP binding sites in the vicinity or even spanning splice junctions.

### The RCRUNCH model for the detection of RBP binding regions

Genome/transcriptome-wide identification of peaks corresponding to individual binding sites for an RBP is time-consuming. For this reason RCRUNCH implements a two-step process, as previously done for analyzing chromatin immunoprecipitation data [14]. That is, broader genomic regions that are enriched in reads in the foreground (IP) sample compared to the background are first identified, and then individual peaks are fitted to the IP read coverage profiles within the selected windows. More specifically, we tabulate the number of fragments that map to sliding windows of a specific size (e.g. 300 nucleotides, sliding by 150 nucleotides at a time), the same in both foreground (IP) and background (e.g. SMI) samples. For windows that are not enriched in binding, fluctuations in the number of reads across replicate samples have been found to be well-described by the convolution of a log-normal distribution due to multiplicative noise in the sample preparation and Poisson sampling noise [14]. The frequency of reads in windows with no RBP binding should in principle be the same between the foreground and the background sample. However, as in the foreground sample reads are expected to come largely from bound regions, the unbound regions will be somewhat depleted of reads. Including a correction term *µ* to account for this depletion, the probability of the data for an unbound region can be modeled as

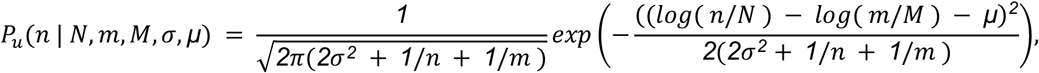

where *n* is the number of reads in a given window (of a total of *N* reads) in the foreground sample, *m* is the number of reads (of a total of *M*) in the background sample, *2σ*^*2*^ is the variance due to multiplicative noise in the two samples, and *1*/*n* and *1*/*m* are the variances due to the Poisson noise. For the probability of read counts due to binding, we assume a uniform distribution over the range corresponding to the maximum and minimum difference in read frequency between foreground and background samples across all windows:

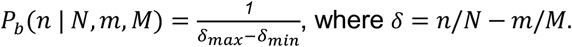

Finally, the probability of observing the *n* reads is given by a mixture model, of the window representing a background region (with probability *ρ*) or a region of RBP binding (with probability *1* − *ρ*):

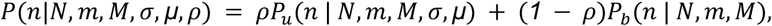

We fit the parameters *ρ, σ, µ* by expectation-maximization (detailed method described in [14], Supplemental material, section 1.9). Regions that are found to be enriched in the foreground sample (z-score higher than 2) are analyzed individually with a second mixture model, to fit peaks corresponding to the individual binding events. The maximum number of peaks considered in a region is the region length divided by the fragment size. The z-scores of these peaks are recalculated based on the number of fragments that are associated with the peaks and those that still have a high enough z-score (corresponding to FDR <= 0.1 for the enriched regions) are kept as significant. When peaks overlap by more than half of the size of one of the peaks, the least significant peak is removed, and the expectation-maximization is continued from the current peak configuration. The procedure is repeated a number of iterations which is twice the maximum number of considered peaks and the configuration with the maximum likelihood score is kept.

### *De novo* motif identification and enrichment calculation

For each peak we extract the region covering 20 nucleotides (nts) upstream and 20 nts downstream of the peak center in the case of the RCRUNCH peak center approach (option ‘peak_center’ in the config). For RCRUNCH crosslink we extract the same type of window, centered not on the peak center but rather on the position where most read starts within the peak are located. To avoid double-counting of motifs, we merge overlapping peaks, obtaining thus a set of non-redundant peaks. To estimate the enrichment of known and *de novo* motifs we use the MotEvo software [27] as described in [14]. Namely, a subset of the peaks is used to train a prior probability of non-specific binding, and the motif enrichment is then estimated from the test set, using this prior. We carried out this procedure 5 times (parameter ‘runs’, can be modified by the user), to both estimate the mean enrichment for known motifs from the ATtRACT database [23], and to identify *de novo* motifs (represented as positional weight matrices, PWMs) of various sizes (provided by the user, default values: 6, 10, 14) that are enriched in the foreground peaks relative to unbound sequences using PhyloGibbs [64]. As unbound sequences we use those genomic regions obtained in the IP experiment that had the lowest z-scores, meaning that they were depleted in IP reads. We sampled 20 background sequences of equal size (40 nts) for each foreground peak. For each motif length, we extracted the top two motifs in the order of cross-validated enrichment and trimmed off positions from the boundaries of the motif until the information score became at least 0.5. For both the known motifs, as well as the *de novo* motifs we report the motif and corresponding enrichment for each of the 5 runs, as the motif can vary to some extent from run to run. The *de novo* motifs from all these runs are then collected together in an extra step and along with the known motifs, the enrichment over all the significant peaks is estimated (in this step there cannot be any prior estimate).

### Benchmarking peak finder tools

We compared RCRUNCH with the recently developed and broadly used tools *CLIPper* [22], *Pure-CLIP* [18] and *omniCLIP* [15]. To reliably and reproducibly perform this analysis we created a separate snakemake workflow. For *PureClip* and *omniCLIP*, docker images [65] were either created or used from existing repositories. As we could not implement *CLIPper* in the same type of workflow as the other tools, we relied on the bed files of *CLIPper-*predicted binding sites from each sample provided by ENCODE. To benchmark the tools we used eCLIP data generated for some RBPs whose binding motifs are well-known: hnRNPC [66,67], IGF2BP3 [66,68], PTBP1 [69,70], PUM2 ([71]) and RBFOX2 [70,72,73]. For *PureClip* and *omniCLIP* the eCLIP data had to be pre-processed separately, as the tools use alignments as input. To facilitate the comparison across methods, we used the pre-processed data from RCRUNCH. The execution of RCRUNCH was done using some specific options as explained in the RCRUNCH workflow description. Firstly, only unique mappers were aligned to the genome. PCR deduplication was performed, using the UMIs that eCLIP experiments contain. We did not remove any reads mapping to ncRNAs, and we used the RCRUNCH genomic approach. Each of the different peak calling methods was applied, and the top 1000 peaks were extracted for the motif analysis, irrespective of FDR threshold. The motif analyses are included as a post-processing part of RCRUNCH. For RCRUNCH, both the ‘crosslink’ and ‘peak center’ positions were used as anchors for extending by 20 nts on either side and obtaining the sequences for the motif analysis. For the other methods, we used the method-predicted crosslink positions as anchors for extracting similar regions of 40 nts in length. For all methods overlapping peaks were merged to ensure non-redundancy in the sequence set. To ensure comparability, we used the same set of sequences with lowest z-scores as background for motif enrichment estimation in the peaks predicted by all samples.

### Calculation of peak agreement between replicate samples and between methods

To calculate the peak agreement between methods and across replicates we used the jaccard distance metric, defined as:

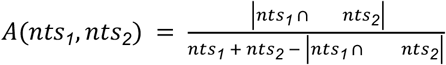, where *nts*_1_ and *nts*_2_ are the total number of nucleotides contained in the top number of peaks chosen for sample 1 and sample 2, respectively.

### Calculation of motif similarity

We defined the motif similarity M of two sequence motifs *m*_1_ and *m*_2_, 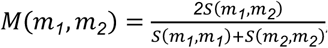, where *S*(*m*_*1*_, *m*_*2*_) = *max*_1_[*I*(*m*_*1*_, *m*_*2*_, *d*)] and *I*(*m*_*1*_, *m*_*2*_, *d*) = *Σ*_2_*m*_*1*_(*i*) *m*_*2*_(*i* − *d*) is the inner product of the motifs with the second motif being at offset *d* compared to the first motif [14]. This measure allows for the comparison of motifs of different lengths, and takes values between 0 (when the base frequency vectors are orthogonal) and 1 (when the two motifs are identical).

### RCRUNCH analysis of ENCODE eCLIP data

We applied the RCRUNCH workflow to all of the ENCODE eCLIP datasets, consisting of 220 distinct eCLIP experiments with 143 different RBPs in two cell lines (K562, HepG2). As for the benchmarks, we used reads mapping uniquely to the genome, performed read deduplication based on the UMIs, and did not exclude reads mapping to ncRNAs. To identify the RBP-specific binding motifs, we used the top peaks for each RBP, based on FDR threshold < 0.1. Agreement across replicates and motif agreement with existing knowledge were the main metrics of performance evaluation. For the comparison of peaks across RBPs and the inference of RBP-specific motifs we considered proteins for which we had at least 20 peaks in both replicate samples. We applied PhyloGibbs and MotEvo (each run 5 times with different seeds of the random number generator) and collected all motifs reported to be enriched in at least one of the replicates for the corresponding protein. Finally, we identified the motif with the highest sum of log-likehood across all replicate samples for a given protein. This motif is reported in Supplementary Table 3 and shown in Fig. 6d.

### RCRUNCH variants

To evaluate the RCRUNCH variants, we used the same samples that were used for benchmarking the computational methods. The pre-processing and post-processing steps (motif analysis) were the same as those implemented in the benchmark analysis, but the selection of reads, regions and peaks differed. Specifically, in the RCRUNCH transcriptome approach we first construct a reference transcriptome composed of the most abundant isoform of each gene in the CLIP data, and use these transcripts as ‘pseudo-chromosomes’ in the mapping process. We then map reads in an unspliced manner to both this reference transcriptome and to the genome and retain the mappings with the highest score. In cases when the alignments to transcriptome and genome are very close in score (transcriptome - genome scores >= -3 points), we give precedence to the transcriptome mappings and ignore the genomic ones. We then apply the standard RCRUNCH (see section: The RCRUNCH model for the detection of RBP binding regions). In RCRUNCH multi-mappers we consider not only reads that map uniquely to the genome, but also those that have up to 50 of equally good mappings (see section: Alignment of reads to reference genome). For RCRUNCH RNA-seq background we simply used RNA-seq samples that are provided by ENCODE for the cell lines (K562, HepG2) used for CLIP. These were treated the same as the SMI sample.

## Abbreviations

AAE: Alu antisense element
AS: alignment score
cDNA: complementary DNA
ChIP: chromatin immunoprecipitation
CLIP: crosslinking and immunoprecipitation (CLIP)
DNA: deoxyribonucleic acid
FAIR principles: findable, accessible, interoperable, reusable
FDR: False discovery rate
mRNA: messenger RNA
ncRNA: non-coding RNA
PCR: polymerase chain reaction
PWM: positional weight matrix
RBDs: RNA-binding domains
RBPs: RNA-binding proteins
RIC: RNA-interactome capture
RNA: ribonucleic acid
RNA-seq: RNA sequencing
RNPs: ribonucleoprotein complexes
UMI: unique molecular identifier
rRNA: ribosomal RNA
tRNA: transfer RNA
SMI: size-matched input
snRNA: small nuclear RNA

## Declarations

### Ethics approval and consent to participate

Not applicable

### Consent for publication

Not applicable

### Availability of Data and Materials

Project name: RCRUNCH

Project home page: https://github.com/zavolanlab/RCRUNCH. The code and intermediate results are also available from the zenodo repository, under DOI 10.5281/zenodo.7642474.

Programming language: Python

License: Apache-2.0

### Authors’ contributions

M.Z., M.K. and E.v.N. designed the study. M.K. implemented the workflow. M.K. and M.Z. carried out the data analysis and wrote the manuscript.

## Acknowledgements

This work would not have been possible without the sciCORE Team [46] at the University of Basel. We are thankful for their support regarding the computational infrastructure as well as dedicated time and effort to aid us in this project. Special thanks to Mikhail Pachkov who provided guidance and help during the adaptation of parts of the CRUNCH model. Many thanks to the Zavolan Lab who contributed to this work with numerous pieces of advice, during the initial development. Special thanks to Dominik Burri and Christina J. Herrmann for help with testing during development.

## Supplementary data

**Figure S1.**
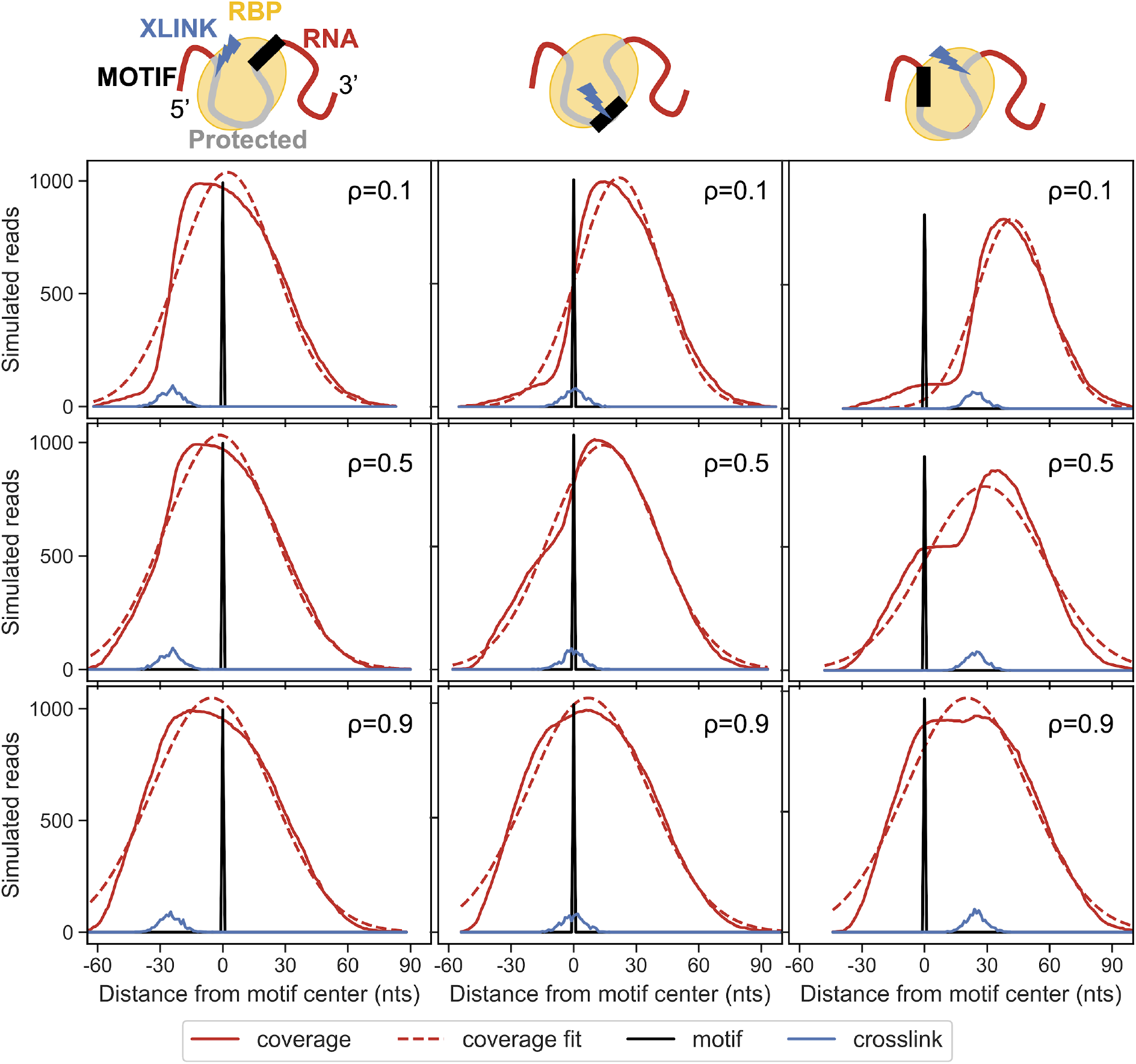
Simulation of a CLIP experiment. Upper schema: an RBP binding to its cognate motif, (black box), crosslinking to the RNA at the site indicated by the blue arrow, and protecting an extended region of the target (shown in gray) from digestion. From right to left, the relative position of the crosslink is changed relative to the motif position. The different columns correspond, as the schema shows, to different positions of the crosslink relative to the motif position. Each row corresponds to different probabilities of readthrough, varying from lower to higher from top to bottom.

**Figure S2.**
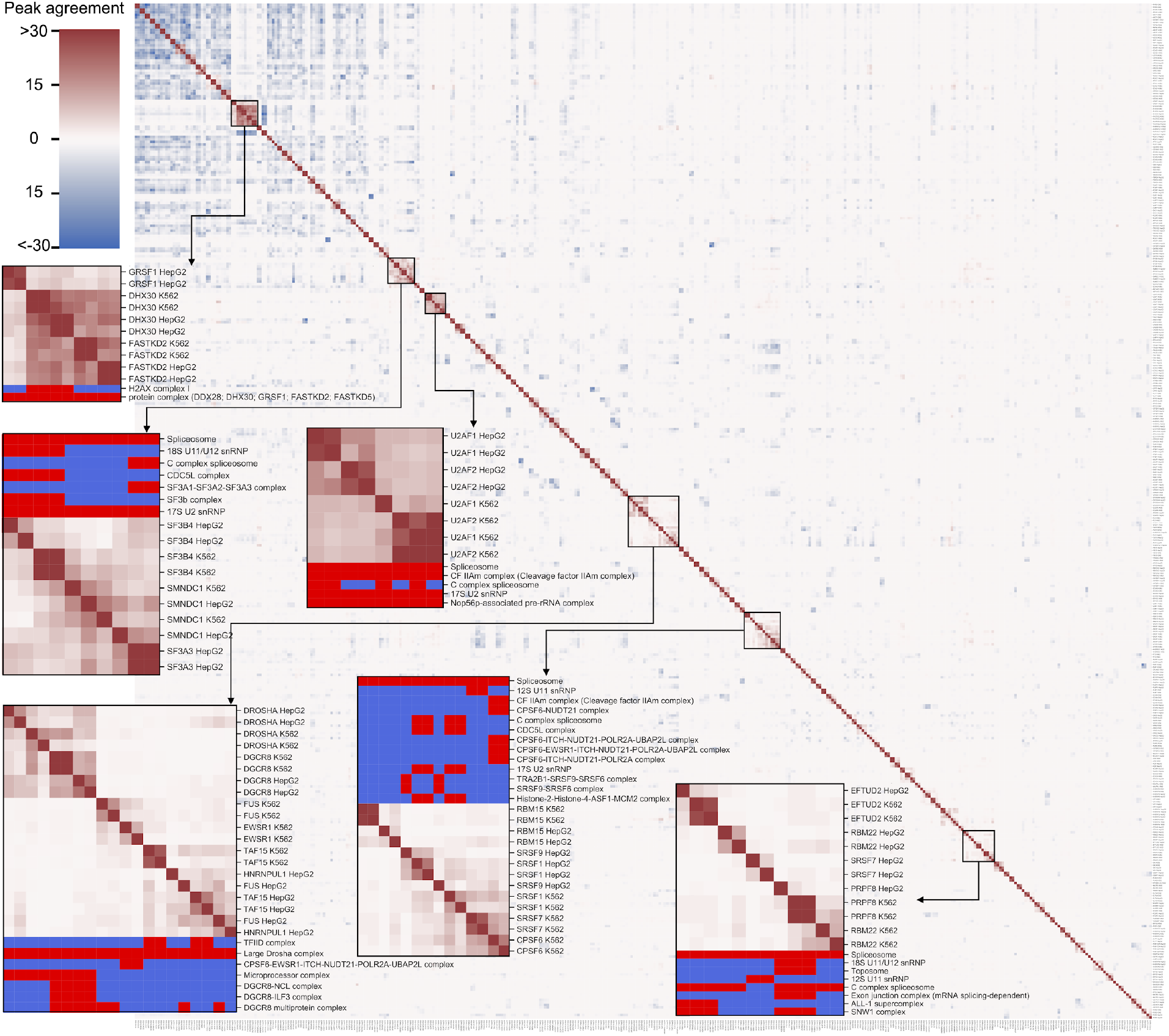
Agreements of peaks identified between individual ENCODE samples. The agreement is calculated as the Jaccard distance of the nucleotides in the peaks, where the intersection of two sets of peaks is the number of nts covered in both sets, while the union is the number of nts covered in at least one of the two sets. The color range is capped at a similarity of 0.4 to make the clusters more easily distinguishable. The top peaks are taken according to the FDR threshold (0.1), extending by 20 nts upstream and downstream from the crosslink site. Only samples with more than 100 peaks are included in this plot. The membership of RBPs in complexes is taken into account (based on CORUM [74]), by multiplying the value of the agreement by 1 if two proteins are known to participate in the same complex, and -1 otherwise. Resulting negative values are shown in blue, while positive values are shown in red. That is, shown in red are peak agreements of samples that either correspond to the same protein, or to proteins known to interact with each other in complexes. On the right, a few clusters of samples containing proteins that are known to take part in complexes are highlighted.

**Figure S3.**
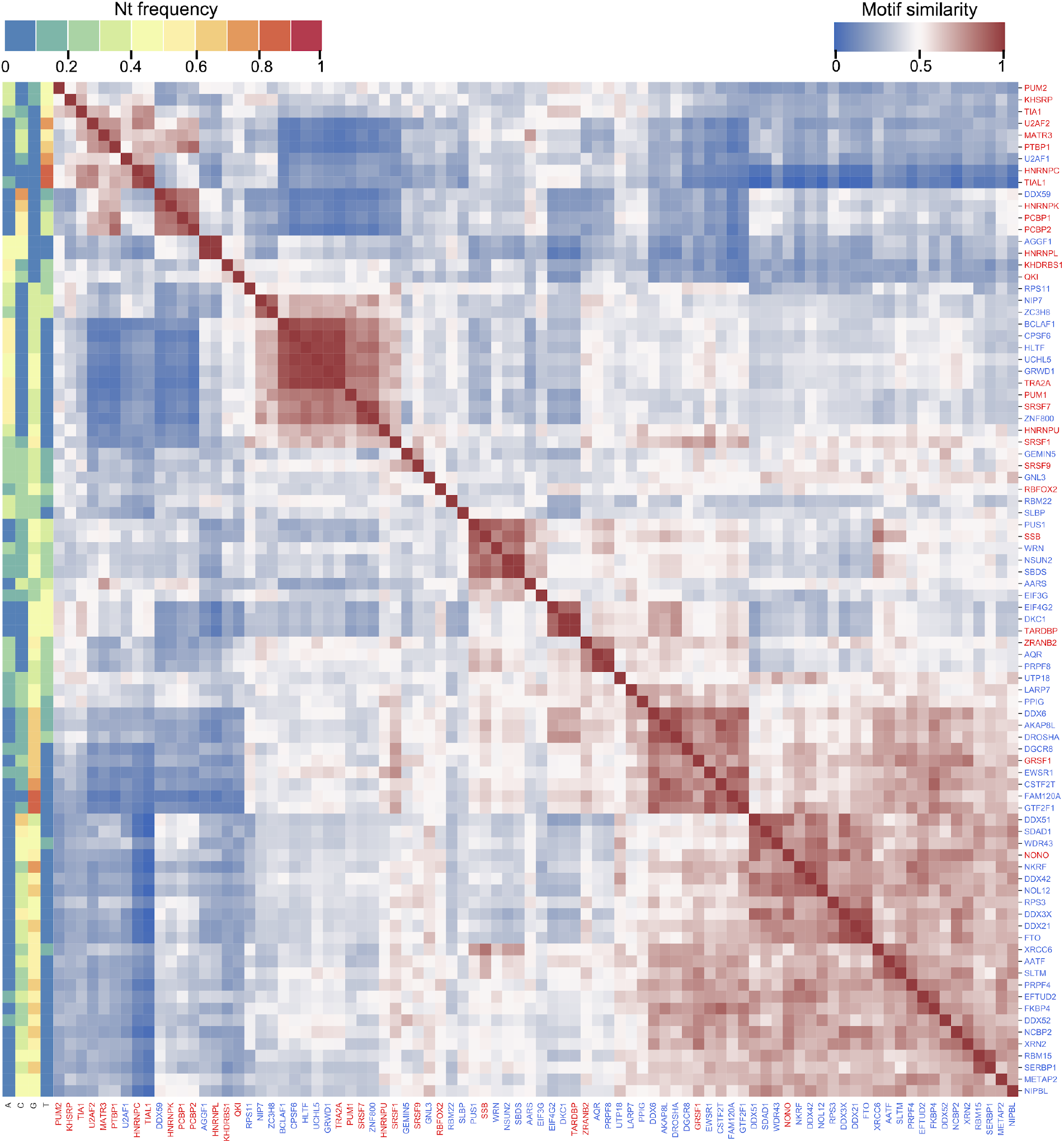
Similarity of *de novo* predicted motifs of different RBPs. Clustermap of *de novo* motif similarity of all RBPs covered in the ENCODE dataset. Similarity is calculated as described in the methods section (Calculation of motif similarity), taking for each RBP the motif that best explains all of the samples corresponding to that RBP. On the left, a representation of the nucleotide composition of the motif is shown, each column corresponding to one nucleotide, while the color indicates the relative frequency of that nucleotide averaged over all positions of the PWM. The RBP names are colored according to whether they have a motif (red) or not (blue) in ATtRACT.

**Figure S4.**
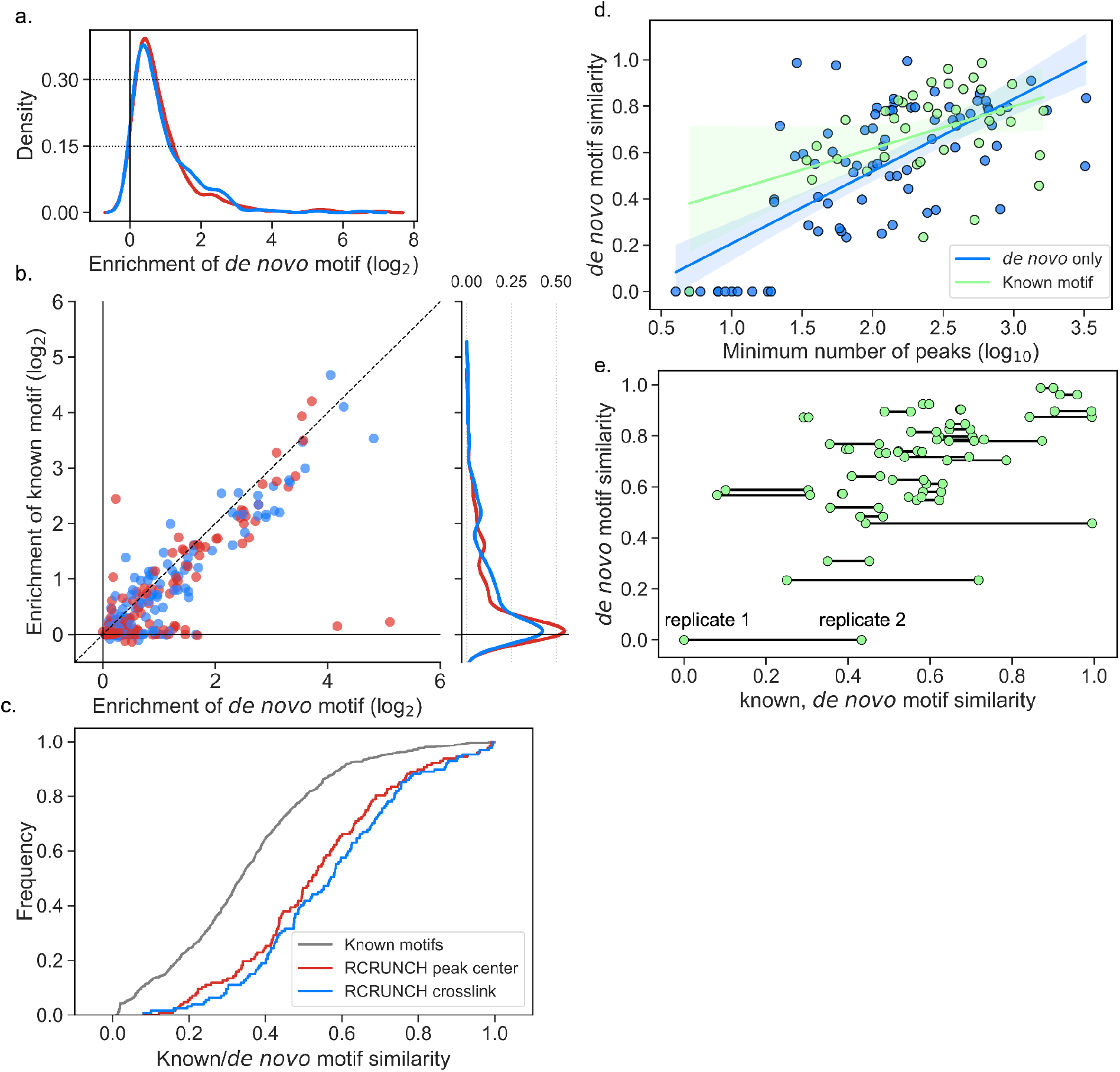
RCRUNCH results for all ENCODE eCLIP data currently available. **a**. Distribution of the enrichment scores of *de novo* motifs computed from peaks identified by RCRUNCH peak center (red) or RCRUNCH crosslink (blue). **b**. Scatter plot of the enrichment of the *de novo* motif predicted in the RCRUNCH-identified peaks from each sample, versus the enrichment score of the known motif for the protein assayed in the respective experiment. Marginal distributions of the known motif enrichments in RCRUNCH crosslink (blue) and RCRUNCH peak center (red) peaks are also shown. **c**. Cumulative density function of pairwise similarity scores for random pairs of known RBP-binding motifs (gray), known and *de novo* motifs identified from RCRUNCH crosslink sites (blue), known and de novo motifs identified for RCRUNCH peak center sites of individual RBPs (red). The same known motif was used for a given protein. **d**. Relationship between the similarity of *de novo* motifs inferred from replicate experiments and the minimum number of binding sites identified in these replicates. Experiments (each corresponding to an RBP and cell line) are colored according to whether (green) or not (blue) a motif was found in ATtRACT for the assayed RBP. **e**. Relationship between the similarity of *de novo* motifs identified in replicate experiments for a given RBP and the similarities of these *de novo* motifs and the known motif of the corresponding RBP. Lines connect replicate samples.

**Figure S5.**
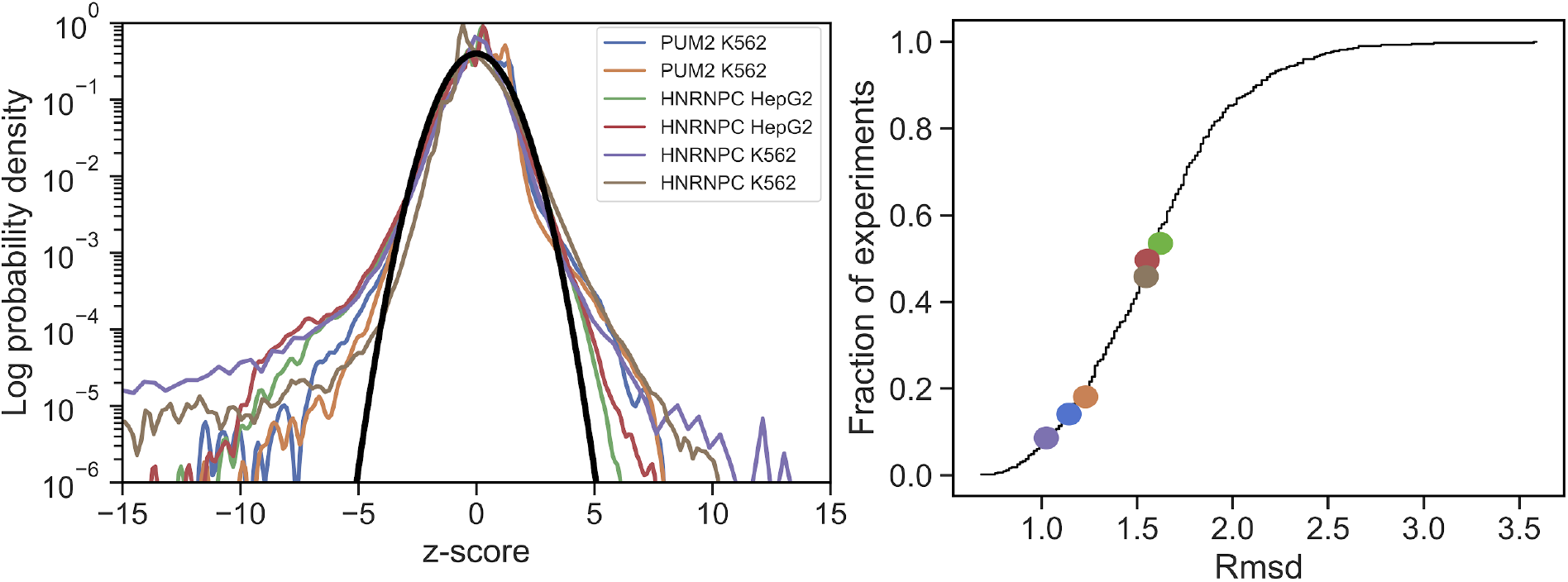
Evaluation of the model used to identify enriched regions. Left panel: z-score distributions for all the genomic windows with at least 2 reads in the IP samples for a random subset of samples (of benchmarked RBPs), compared to the expected distribution (in black). Right panel: cumulative distribution of root-square mean deviation of the empirical and predicted z-score distribution for all eCLIP samples in ENCODE. The samples shown in the left panel are highlighted with the same colors.

**Figure S6.**
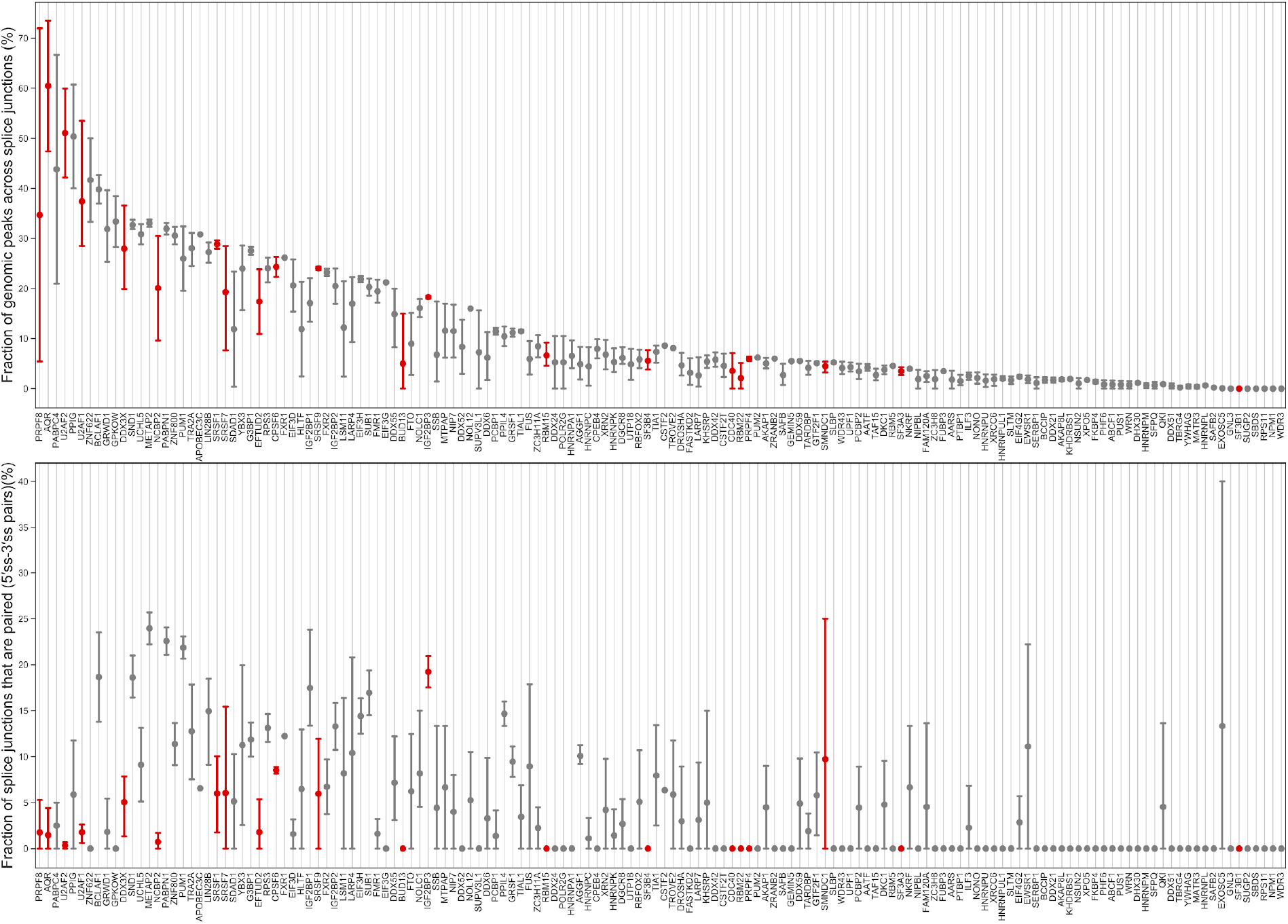
Binding events spanning splice junctions. **a**. Scatterplot of average fraction (and standard deviation) of peaks identified for the specified RBPs by RCRUNCH (genomic approach) that overlap with annotated splice junctions. The samples are ENCODE eCLIP samples and the average is calculated over replicates and cell lines. **b**. Scatterplot showing the fraction of the peaks that overlap a junction (e.g 3’ss) and also have a “pair” peak which overlaps the other side of the same junction (5’ss).

**Figure S7.**
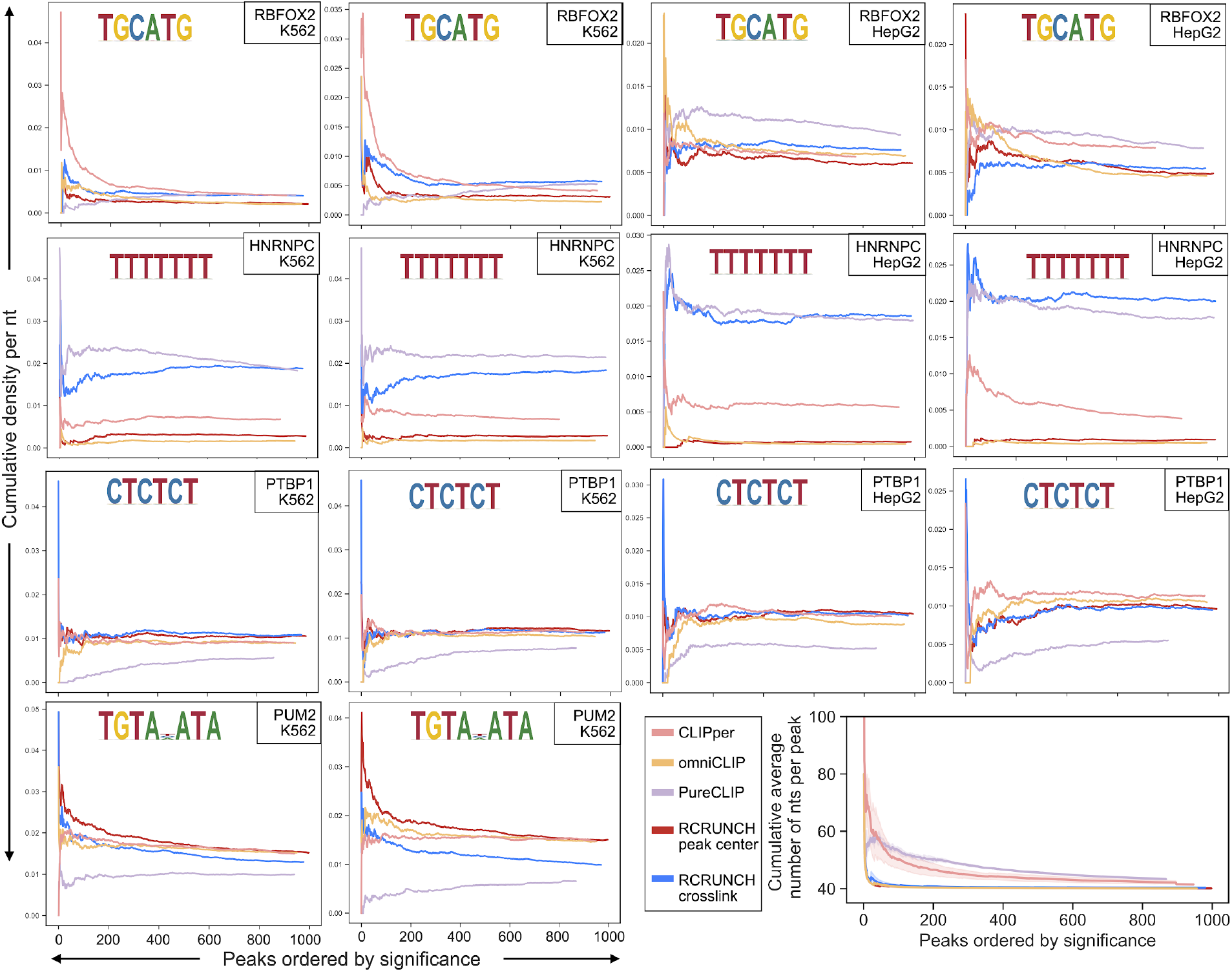
Motif enrichments as a function of the number of top peaks. Analyses were carried out for individual samples (IP vs. corresponding SMI) corresponding to one RBP and cell line. The top *x* peaks identified by a given tool, according to the tool’s significance measure, were used to calculate posterior probabilities of sites (matching known motifs from ATtRACT) with the MotEvo tool [27] assuming uniform nucleotide frequencies and a prior probability for the background of 0.99. Sites with a posterior >= 0.3 per nucleotide in the peaks were used to calculate the average site density per nucleotide in peaks. The figures show that in general, the top peaks have the highest density of expected motifs. Bottom right panel: average number of nucleotides per peak as a function of the number of top peaks, sorted by significance. For each tool (indicated by the color), peaks were extracted from a given sample, overlapping peaks were merged and average of merged peak sizes over all benchmarked proteins and all samples were calculated. The standard deviations are also shown. The figure shows that CLIPper and PureCLIP tend to extract closely-spaced peaks, which are then merged to yield broader peaks.

**Figure S8.**
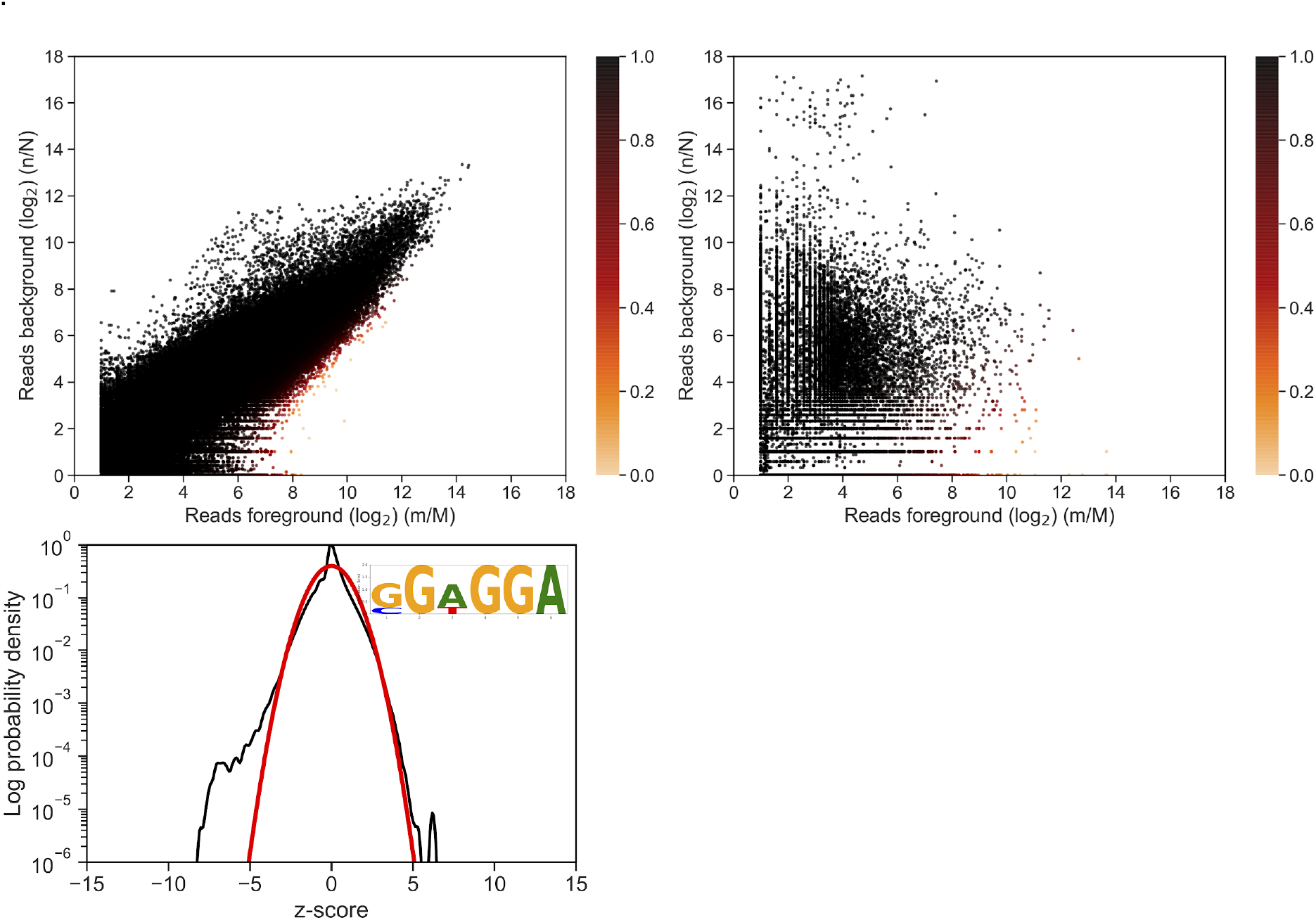
RCRUNCH application to PAR-CLIP data. RCRUNCH analysis was performed on two example datasets: for the fly CNBP protein [75], known to bind a GGA motif [15,75], and the human PUM2 protein [76], known to bind the UGUANAUA motif. The CNBP IP sample had a matched SMI control, whereas for the PUM2 PAR-CLIP we used as control RNA-seq data from [76,77].

